# The pseudouridine synthase dyskerin binds to cytoplasmic H/ACA-box snoRNA retaining transcripts affecting nuclear hormone receptor dependence

**DOI:** 10.1101/2021.01.21.427585

**Authors:** Federico Zacchini, Giulia Venturi, Veronica De Sanctis, Roberto Bertorelli, Claudio Ceccarelli, Donatella Santini, Mario Taffurelli, Marianna Penzo, Davide Treré, Alberto Inga, Erik Dassi, Lorenzo Montanaro

**Affiliations:** Dipartimento di Medicina Specialistica, Diagnostica e Sperimentale (DIMES), Alma Mater Studiorum – Università di Bologna, Bologna, I-40138 - Italia; Centro di Ricerca Biomedica Applicata - CRBA, Università di Bologna, Policlinico di Sant’Orsola, Bologna, I-40138 - Italia; Dipartimento di Biologia Cellulare, Computazionale e Integrata (CIBIO), Università di Trento, I-38123 Trento - Italia; Unità Operativa di Anatomia Patologica, Azienda Ospedaliero-Universitaria di Bologna, Via Albertoni 15, I-40138 Bologna - Italia; Unità Operativa di Chirurgia Generale, Azienda Ospedaliero-Universitaria di Bologna, Via Albertoni 15, Bologna I-40138 - Italia; Dipartimento di Scienze Mediche e Chirurgiche (DIMEC), Alma Mater Studiorum - Università di Bologna, Bologna, I-40138 - Italia; Programma Dipartimentale di Medicina di Laboratorio, Azienda Ospedaliero-Universitaria di Bologna, Via Albertoni 15, Bologna, I-40138 - Italia

**Keywords:** DKC1, intron retention, RNA binding, post-transcriptional control, breast cancer

## Abstract

Dyskerin is a nuclear protein involved in H/ACA box snoRNA-guided uridine modification of RNA. Since its defective function induces specific alterations in gene expression, we sought to unbiasedly identify mRNAs regulated by dyskerin. We found that dyskerin depletion affects the expression or the association with polysomes of selected mRNA isoforms characterized by the retention of H/ACA box snoRNA-containing introns. These snoRNA retaining transcripts (snoRTs) are bound by dyskerin and can interact with cytoplasmic ribosomes. We then characterized the cytoplasmic dyskerin RNA interactome finding both H/ACA box snoRTs and protein-coding transcripts. Since a fraction of these latter transcripts is involved in the nuclear hormone receptor binding, we tested to see if this specific activity is affected by dyskerin. Results indicate that dyskerin dysregulation may alter the dependence on nuclear hormone receptor ligands in breast cancer. Our work suggests a cytoplasmic function for dyskerin which could affect mRNA post-transcriptional networks relevant for nuclear hormone receptor functions.

## INTRODUCTION

Dyskerin is a conserved, multifunctional protein encoded by the DKC1 gene (1). DKC1 mutations cause the rare multisystemic syndrome X-linked dyskeratosis congenita (1). In addition, dyskerin expression is often dysregulated in human cancer (2).

Major dyskerin functions include telomerase complex stabilization (3) and the site-specific isomerization of uridine to pseudouridine in RNA molecules (4). These functions are achieved by dyskerin binding to a class of non-coding small nucleolar RNAs termed H/ACA box snoRNAs, which also include the telomerase RNA component (TERC). Dyskerin binds these RNA molecules in association with other pseudouridylation complex core proteins, namely NHP2, NOP10, and GAR1(5,6).

On the basis of their specific sequence, most H/ACA box snoRNAs guide the pseudouridylation complex on specific uridine residues for their modification to pseudouridine. The majority of these target uridines lie on ribosomal RNA (rRNA) (7).

In vertebrates, most of H/ACA box snoRNA genes are present as intronic sequences contained in a subset of essential genes involved in the synthesis or functioning of the translational apparatus, including those coding for ribosomal proteins, translation factors, nucleolar proteins, and proteins involved in mRNA binding, transport and stability. H/ACA box snoRNAs are then transcribed by RNA Pol II and generated mainly through the splicing of the nascent pre-mRNA and the exonucleolytic trimming of the spliced intron (8–10).

Pathogenic DKC1 mutations and dyskerin depletion induce both TERC destabilization with a loss of telomerase activity (3) and a defective rRNA pseudouridylation (2,11–13), associated with changes in mRNA translation which end up with specific alterations in gene expression. Such changes involve altered cap-independent translation initiation and reduced translational fidelity (14–18). Notably, these effects were involved to explain the role played by dyskerin in the development of different tumour types, including breast cancer(11,16,17,19).

In this study, we sought to identify mRNAs affected by dyskerin depletion at the transcriptome-wide level. We thus performed an RNA-seq analysis of total and polysome-associated RNAs in control and stably dyskerin partially depleted breast cancer-derived cells. Results showed that DKC1 mRNA downregulation strongly affects the recruitment to polysomal fractions of mRNA isoforms characterized by the retention of H/ACA box snoRNA sequences containing introns. Further analyses indicated that dyskerin binds to these snoRNA retaining transcripts (snoRTs) in the cytoplasm and is involved in the regulation of a complex RNA interactome affecting mRNA post-transcriptional networks, which are important for nuclear hormone receptor functions.

## MATERIAL AND METHODS

### Cell Culture

MCF-7 (female, oestrogen-positive invasive breast ductal carcinoma derived) cells were cultured in RPMI 1640 medium supplemented with 10% FBS, 100 U/ml penicillin, 1 mg/ml streptomycin, and 2 mM L-glutamine. MDA-MB 231 (female, triple-negative invasive breast ductal carcinoma-derived) cells were cultured in DMEM containing 10% FBS, 1 mg/ml streptomycin, and 2 mM L-glutamine. Cells were maintained at 37°C and 5% CO_2_. Cell lines were obtained from the American Type Culture Collection (ATCC) and were routinely tested for mycoplasma using Venor®GeM Classic (Minerva Biolabs). Stable cell lines were generated by transfecting MCF7 cells with a retroviral vector expressing a short hairpin RNA (shRNA) targeting DKC1, as described previously (17). The sequence of the short hairpin oligonucleotide targeting DKC1 mRNA was 5’-CCAAGGTGACTGGTTGTTTAAT-3’. As a negative control, an empty vector was used. Stable retroviral-transduced populations were selected in standard medium supplemented with 4 μg/ml blasticidin. For siRNA-mediated depletions, cells were transfected with siRNAs targeting DKC1 (Invitrogen, catalogue number HSS102781: 5’-AACACCUGGAAGCAUAAUCUUGGCC-3’, HSS102782: 5’-UAAACAACCAGUCACCUUGGGAUCC-3’, HSS102785: 5’-GAAGUCACAACAGAGUGCAGGCAAA-3’) and an appropriate control (Cat. N° 12935300 - Invitrogen) using Lipofectamine RNAiMAX reagent (Invitrogen) in OptiMEM medium (Invitrogen) according to the manufacturer’s instructions. Cells were harvested 72 h after the transfection. For experiments that required the removal of ligands of nuclear hormone receptors, MCF7 cells with stable dyskerin depletion and the relevant control were cultured in phenol red-free medium containing 10% charcoal-treated FBS for four days.

### Cells lysis and subcellular fractionation

MCF7 and MDA-MB 231 cells were lysed with different buffers: for total protein recovery (for western blot analysis) cells were lysed in RIPA buffer (50 mM Tris-HCl pH 7.5, 150 mM NaCl, 1% IGEPAL, 0.1% and protease inhibitor cocktail) according to a standard procedure. For the RNA Immunoprecipitation assay from the whole cellular lysate, cells were washed in phosphate-buffer saline, collected by scraping, and dissolved in Immunoprecipitation Buffer (25 mM Tris-HCl pH 7.5, 150 mM KCl, 5 mM MgCl_2_, 1mM EGTA, 10% glycerol, 1.5% IGEPAL, 0.05% SDS and protease and RNAse inhibitor cocktail) for 30 min at 4°C, followed by centrifugation at 12000 g for 20 min at 4°C. The supernatant was collected for the total RNA IP assay. For subcellular nuclear/cytoplasmic fractioning (for the RIP and polysome assays), cells were washed in phosphate-buffer saline, collected by scraping, and dissolved in Cytoplasm Lysis Buffer (15 mM Tris-HCl pH 7.5, 7.5 mM NaCl, 1.5 mM MgCl_2_, 0.3% IGEPAL, 50 mM sucrose and protease, and RNAse inhibitor cocktail) for 15 min, followed by centrifugation at 1200 g for 15 min at 4°C. The supernatant was collected as the cytoplasmic fraction and, if needed, the pellet was washed in PBS (followed by centrifugation at 1000 g for 10 min at 4°C) and then resuspended in Nucleus Lysis Buffer (50 mM Tris-HCl pH 7.5, 50 mM KCl, 300 mM NaCl, 10% glycerol, 0.1 mM EDTA, 0.5% IGEPAL, and protease and RNAse inhibitor cocktail) for nuclear lysis, and finally centrifugated at 10000 g for 3 min at 4°C. The supernatant was collected as nuclear fraction.

### RNA Immunoprecipitation (RIP)

For RIP assay, a total amount of lysate corresponding to 500 μg of protein content from MCF7 and MDA-MB 231 cells were pre-cleaned with protein A/G Plus-Agarose beads (Santa-Cruz, sc-2003) for 1 hour at 4°C. Five to ten percent of the pre-cleaned sample was saved as input for subsequent analysis. The remainder was used in immunoprecipitation reactions with rabbit IgG (Santa Cruz, sc-2027) or rabbit polyclonal antibody against dyskerin (Genetex GTX 109000) with incubation overnight at 4°C. Afterwards, an anti-dyskerin interacting fraction was captured using protein A/G Plus-Agarose beads and washed several times with Wash Buffer (25 mM Tris-HCl pH 7.5, 150 mM KCl, 5 mM MgCl_2_, 1mM EGTA, 10% glycerol and protease and RNAse inhibitor cocktail). One third of the immunoprecipitated solution was saved for Western blot analysis, while the remainder was used for RNA extraction. Likewise, after saving inputs of cytoplasmic/nuclear cell lysates, the samples were entirely subjected to the same RIP analysis.

### RNA isolation and reverse transcription

Total RNA was purified using the PureZOL™ RNA Isolation Reagent (BIORAD) according to the manufacturer’s guideline. cDNA was obtained retrotranscribing 500 ng of RNA using an iScript™ cDNA Synthesis Kit (BIORAD) according to the manufacturer’s guideline. The RNA derived from polysome isolation and RIP purification (and respective control/input) were purified using the standard phenol:chloroform approach, briefly: the proteins were digested adding proteinase K (final concentration of 100 μg/mL) and SDS (final concentration of 1%) at 37°C for 1h. Afterwards, the RNA was extracted with 1/4 volume of phenol:chloroform 5:1 acid equilibrated (pH 4.7) and NaCl (final concentration of 500 mM) by spinning at 16000 g for 5 min at 4°C. The RNA was then precipitated from the aqueous phase using isopropanol. All the RNA obtained by purification was retrotranscribed using an iScript™ cDNA Synthesis Kit (BIORAD) according to the manufacturer’s guideline.

### Quantitative PCR

qPCR analyses were performed using Taqman probes (SsoAdvanced Universal Probes Supermix, BIORAD) or SYBR green (SsoAdvanced™ Universal SYBR® Green Supermix, BIORAD). Analyses were conducted via the CFX Connect Real-Time detection System (BIORAD) and the expression level was determined by using CFX Manager Software (BIORAD). The relative expression of different transcripts was normalized to β-actin mRNA or β-glucuronidase (GUS) mRNA as endogenous controls. qPCR primer sequences are listed in Supplementary Table S1.

### Digital PCR

Digital PCR was performed using the QuantStudio 3D PCR system (Applied Biosystem) according to the manufacturer’s instructions. Reactions were prepared using specific primers and probes for the target detection as reported in Supplementary Table S1. The equivalent amount of 100 ng of retrotranscribed RNA was used for the absolute quantification of SNORA67-EIF4A1 species and H/ACA EIF4A1 mRNA; results were normalized against the GUS housekeeping gene.

### Western blotting

Protein solutions were diluted in Laemmli loading dye (2% SDS, 8% glycerol, 62.5 mM Tris-HCl pH 6.8, 0.005% bromophenol blue, and 2% β-mercaptoethanol) and separated on a 10% gel by SDS-PAGE (TGX Stain-Free™ FastCast™ Acrylamide Solutions; BIORAD) or 4-15% gel (Mini-PROTEAN® TGX Stain-Free™; BIORAD). Proteins were transferred to nitrocellulose membranes (Amersham Protran 0.2um nc) and probed overnight at 4°C with appropriate antibodies. Detection was performed with appropriate HRP-conjugated secondary antibodies and chemiluminescent detection was performed on a ChemiDoc XRS+ Imaging System (BIORAD) using ImageLab v5.1.1 (BIORAD). The Stain-Free system makes it possible for us to check the quality and quantity of the loaded proteins, especially for RIP and polysome protein gels. The following antibodies were used: anti-Dyskerin (Santa Cruz Biotechnology, sc-373956), anti-NHP2 (Santa Cruz Biotechnology, sc-398430), anti-NOP10 (Cusabio, CSB-PA873610LD01HU), anti-GAR1 (Proteintech, 11711-1-AP), anti-RPS14 (Santa Cruz Biotechnology, sc-68873), anti-RPL5 (Bethyl, A303-933A), anti-EIF4GI (Cell Signaling, 2858S), anti-Vimentin (Cell Signaling, 5741S), anti-GAPDH (Sigma-Aldrich, G8795), anti-b-Actin (Sigma-Aldrich, A2228), Lamin B (Santa Cruz Biotechnology, sc-6216).

### Polysome fractionation

For the standard polysome fractioning, cultured MCF7 cells were treated with 100 μg/mL cycloheximide (CHX), while for testing whether transcripts are truly associated with polysomes, cultured MCF7 cells were treated with 100 μg/ml puromycin. Cells were incubated for 20 min at 37°C and washed on ice twice with cold PBS containing 100 μg/mg CHX or 100 μg/ml puromycin. Cells were scraped and lysed in Cytoplasm Lysis Buffer for cytoplasmic sub-fractioning (as described above) with the addictions of CHX or puromycin. The lysate was centrifuged at 14000 g for 10 min at 4°C. 700 μg to 1000 μg of total proteins were layered onto a chilled sucrose gradient (10-50% or 15-30%) in low-salt buffer (LSB) (20 mM Tris-HCl pH 7.5, 10 mM NaCl, 3 mM MgCl_2_, 0.4% IGEPAL, 50 mM sucrose, and RNAse inhibitors) with 100 μg/mg CHX. Gradients were centrifuged at 36000 RPM for 2 hours at 4°C in a SW41 rotor (Beckman Coulter) and then fractions were collected using a gradient collector (Teledyne ISCO gradient station) coupled with UV detector to continue monitoring the absorbance at 254 nm. For mRNA sequencing, 10% of each fraction was pooled to reconstitute the total mRNA, while the remaining polysomal fractions were pooled together. For standard polysome profiling, one half of each fraction was used for protein purification and the other half for RNA purification. Proteins were recovered using acetone precipitation: the same volume of 100% icecold acetone and 1/10^th^ of TCA were added to each fraction. The samples were placed at −80°C overnight and then centrifuged at 16000 g for 10 min at 4°C. The resulting pellets were washed three times with 1 ml of 100% ice-cold acetone in order to remove TCA residuals and dried at RT for 5 min. Pellets were finally resuspended in a Laemmli loading dye. RNA was purified as described above.

### Ribosome purification

Human ribosomes from MCF7 and MDA-MB 231 cells were purified by lysing cell pellets via the addition of 2 packed cell volumes of 10 mM Tris - HCl, pH 7.5, 10 mM NaCl, 3 mM MgCl_2_, and 0.5% IGEPAL for 10 min at 4°C. The lysate was centrifuged at 20000 g for 10 min at 4°C to isolate the cytoplasmic fraction from nuclei and mitochondria. Highly purified ribosomes (high salts) were obtained as previously described (20). Briefly, 500 μl of lysate were incubated 10 min at 37°C to enable ribosomes to complete translation and become detached from the mRNAs they were translating. Then the cytoplasmic lysate was layered on a discontinuous sucrose gradient in stringent conditions (top-half gradient: 1.0 M sucrose 30 mM Hepes/KOH, pH 7.5, 2 mM magnesium acetate, 1 mM DTT and 70 mM KCl; bottom-half: 0.7 M sucrose, 30 mM Hepes/KOH, pH 7.5, 2 mM magnesium acetate, 1 mM DTT and 0.5 M KCl) by centrifugating samples for 16 h at 160000 g at 4°C. For less stringent purification (low salt), 500 μl of lysate were layered on a single sucrose cushion (1 M sucrose, 10 mM Hepes pH 7.5, 10 mM potassium acetate, 1 mM magnesium acetate, and 1mM DTT) and then centrifugated at the same condition of high salt purification. Ribosomes were washed twice with 10 mM Tris/HCl pH 7.5, 2 mM magnesium acetate, and 100 mM ammonium acetate, and resuspended in the same buffer. Ribosome concentration was calculated from the A260 (1 mg/ml ribosome = 12.5 A260).

### Patients’ material

One hundred twenty breast carcinomas were selected for EIF4A1 mRNA expression determination from a series of consecutive patients who underwent surgical resection for primary breast carcinoma at the Surgical Department of the University of Bologna, on the sole basis of frozen tissue availability. Some of the cases were obtained from a previous study (2). Data on patients’ survival, tumour histological classification, oestrogen and progesterone receptor status were obtained as described elsewhere (2). Informed consent was obtained from all the individual participants included in the study. Data collection was subject to the availability of specific patient information or surgical tissues. All the procedures performed in studies involving human participants meet the ethical standards of the Institutional Research Committee (Nos. 75/2011/U/Tess approved on 7/19/2011 and 132/2015/U/Tess approved on 10/13/2015 by Policlinico S.Orsola-Malpighi Ethical Review Board, Bologna, Italy) and the 1964 Helsinki declaration as subsequently amended or any comparable ethical standards.

### Experiments with Mice

All animal work was approved by Bologna University’s Institutional Animal Care and Use Committee in accordance with national guidelines and standards (protocol approval reference No. 204/2016-PR). Six-week-old female BALB/c nude mice were purchased from Charles River (Charles River Laboratories Italia s.r.l.). The mice were maintained in a specific pathogen-free facility, on a 12-h light-dark cycle at 21°C. To perform the breast cancer xenografts, 5 million stable dyskerin interfered MCF7 cells and the relevant control cell line were injected subcutaneously into both flanks of anesthetized mice (10 mice per cell line). No estradiol supplement was administered to the mice (21). The mice were weighed once a week, and the tumour growth was monitored weekly. The animals were euthanized at post-injection week 33, by an anaesthetic overdose according to the approved experimental protocol.

### mRNA seq

RNA purified from polysomal fractioning (the reconstituted total mRNA and pooled polysomal RNA) was tested for quality using a RNA 6000 nano kit (Agilent) on a Bioanalyzer 2100 (Agilent).

Poly-A enriched, strand-specific RNA libraries were generated with the TruSeq mRNA Stranded sample preparation kit (Illumina) starting from 1μg of RNA from each fraction (total and polysomal). Briefly, RNA was subjected to poly(A) selection using Magnetic Oligo-dT Beads. Poly(A^+^) RNA was partially degraded by incubating in Fragmentation Buffer at 94°C for 4 min. After first- and second-strand cDNA synthesis using random primer, end repair and A-tailing modification, Illumina Truseq sequencing adaptors were ligated to cDNA ends. cDNAs were amplified by 15 cycles of PCR reactions and subsequently purified by AMpure XP beads (Beckman). Each individual library was quantified, and quality controlled using a Qubit Fluorometer (Thermo Scientific), LabChip GX (Perkin Elmer). After libraries equimolar pooling, the final pool was quantified by qPCR (KAPA and BIORAD). The adaptor-tagged pool of libraries was loaded on an Illumina Hiseq2500 high throughput flowcell (PE50 chemistry) for cluster generation and deep sequencing. Reads were filtered by quality and trimmed, and adapters removed (minimum quality 30, minimum length 36 nt) with Trimmomatic (22). Reads were then aligned to the hg38 genome with STAR (23), using the Gencode v28 (http://www.gencodegenes.org/releases/) annotation to quantify genes. Transcripts were aligned and quantified with Salmon (24) on the same annotation. DESeq2 (25) was then used to call Differentially Expressed Genes (DEGs) between conditions (shDKC1 vs control) at the total and polysomal level, using a 0.05 threshold on the adjusted p-value (Data are available on Supplementary Table S2).

Exon analysis was performed on aligned reads with DEXseq (26), using an adjusted p-value threshold of 0.05 to compare conditions at the total and polysomal level.

The functional enrichment analysis on Gene Ontology annotations was performed by the topGO R package (27), using a BH-adjusted p-value threshold of 0.05 (Data are available on Supplementary Table S3).

### RIP seq

RIP assay was performed as described above, and the cytoplasmic RNA purified from RIP was tested for quality using a RNA 6000 nano kit (Agilent) on a Bioanalyzer 2100 (Agilent), while 10 ng of the extracted RNA was used for fragmentation at 94°C for 4 min. RNA libraries were generated using the SMART-Seq Stranded Kit (Takara). This kit incorporates SMART® cDNA synthesis technology (28) and generates Illumina-compatible libraries via PCR amplification, thus avoiding the need for adapter ligation and preserving the strand orientation of the original RNA. The Ribosomal cDNA was depleted by a ZapR-mediated process, in which the library fragments originating from rRNA and mitochondrial rRNA are cleaved by ZapR in the presence of mammalian-specific R-Probes. Library fragments originating from non-rRNA molecules were enriched via a second round of PCR amplification using Illumina-specific primers and, subsequentially, purified. Each individual library was quantified and quality-controlled using a Qubit Fluorometer (Thermo Scientific), LabChip GX (Perkin Elmer). Equal numbers of cDNA molecules from each library were pooled and the final pool was purified once more in order to remove any free primer. Following a final qPCR quantification (KAPA and BIORAD), the pool was loaded on a Hiseq2500 rapid run flow-cell and run in a PE50 chemistry. Reads were preprocessed (quality threshold Q30, minimum length 36nts, adapters removed) using TrimGalore (https://www.bioinformatics.babraham.ac.uk/projects/trim_galore/), then aligned to the hg38 genome with STAR (23), using the Gencode v28 (http://www.gencodegenes.org/releases/) annotation to quantify genes. Transcripts were aligned and quantified by Salmon (24) on the same annotation. Gene read counts were normalized by library size. RIP fold-enrichment and p-value were computed for each condition using DESeq2 (25) as (RIP / INPUT) or (IGG / INPUT). Genes and transcripts which became significantly enriched (adjusted p-value <= 0.05) in the RIP/INPUT and not in the corresponding IGG/INPUT were considered to be dyskerin targets. The functional enrichment analysis of dyskerin targets was performed as previously described.

### Statistical analysis

Statistical analyses (tests, number of replicates, and two-sided p-values) are indicated in the corresponding figures or figure Legends.

## RESULTS

### Dyskerin regulates the recruitment to polysomes of snoRTs

We stably reduced the levels of dyskerin in breast carcinoma-derived MCF7 cells by specific shRNA expression as previously performed (17) (**Figure 1A**). These cells, and their relevant controls, were used to isolate total and actively translated cytoplasmic mRNAs by polysomal fractionation. PolyA - RNA-Seq was then performed for an unbiased identification of mRNAs whose translation is affected by dyskerin depletion (**Figure 1B**). First, the results were analysed at the gene level, showing no significant difference in total and polysomal fractions (except for DKC1) after false discovery rate correction (**Figure S1A**). Then the differences involving specific mRNA splicing isoforms were investigated by both isoform-based and exon-based analysis. These analyses led to the identification of a subset of mRNA isoforms and exon sequences which are either differentially represented in total fractions or differentially recruited to polysomes after dyskerin depletion (see **Figure 1C** and **Figure S1B, S1C**).

**Figure 1.**
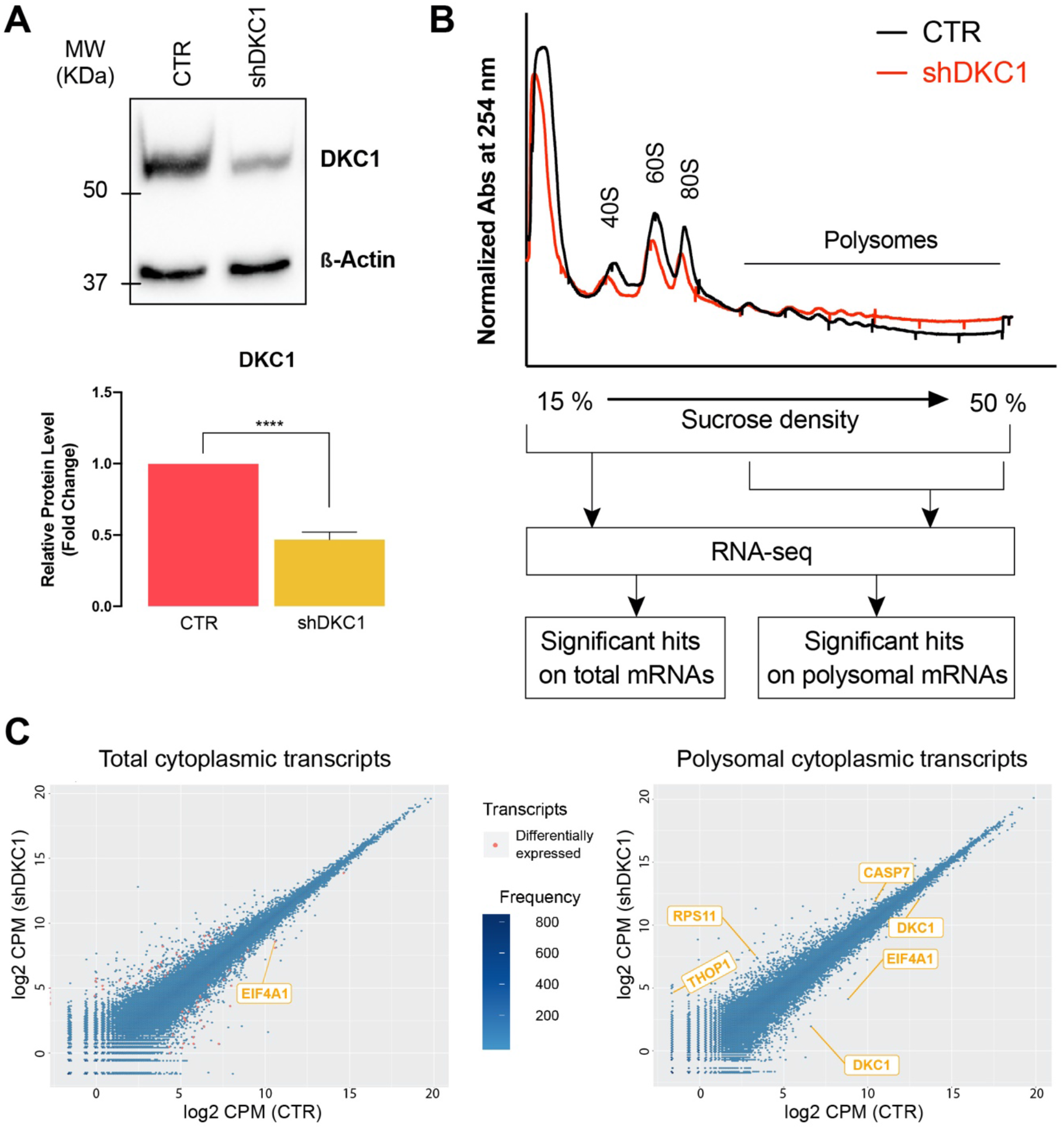
Dyskerin regulates the recruitment to polysomes of snoRT. (A) Representative Western blot analysis image (top) and densitometric analysis of 5 independent replicates (bottom) of dyskerin levels after DKC1 mRNA KD in MCF7 cells. Data are shown as means ± SD. Paired Student’s t tests were performed on controls. ****p < 0.001. (B) Representative polysome profile obtained by 15-50% sucrose density gradient centrifugation from control (black) and DKC1 KD (red) MCF7 cells. Three independent replicates were used. Ten percent of each fraction was pooled to reconstitute total mRNA, while the remaining polysomal fractions were pooled together. Reverse transcribed polysomal and total mRNAs were sequenced using a next-generation sequencing (RNA-seq) approach with a depth of about 50-60 M usable reads. Processed data were analysed for significant differences in mRNA levels normalized signals between polysomal and total RNA fractions. (C) Count-based differential expression analysis of total cytoplasmic (left panel) and polysomal-recruited (right panel) transcripts. Differentially expressed transcripts are depicted as red dots, while transcripts gene names of interest are indicated.

Interestingly, the list included several mRNA isoforms which retain introns containing H/ACA box snoRNA sequences that we defined as H/ACA snoRTs (**Figure 1C** and **Figure S1B, S1C**, highlighted lines). In particular, we identified an isoform of the EIF4A1 mRNA which retains the intron transcribing for SNORA67 (EIF4A1 snoRT), an isoform of RPL32 mRNA retaining SNORA7A (RPL32 snoRT), and isoforms of TAF1D and DKC1 mRNAs retaining several H/ACA box snoRNAs (see **Figure S1B, S1C** for details). The effect of dyskerin depletion on these snoRTs was further validated by RT-qPCR using different siRNA sequences to reduce DKC1 levels and also on a second cell line (**Figure S1D, S1E**). These results, given dyskerin ability to bind to H/ACA box snoRNAs sequences in the nucleus, suggested that H/ACA snoRTs might also be bound by dyskerin.

### H/ACA snoRTs are bound by dyskerin in the cytoplasm

To investigate the association of dyskerin with H/ACA snoRTs, we performed an RNA immunoprecipitation (RIP) using an anti-dyskerin antibody followed by RT-qPCR. For this purpose, we used primers capable of distinguishing the canonical and intron-retaining isoforms of interest (**Figure 2A, Figure S2**). The results obtained on MCF7 and MDA-MB-231 breast cancer-derived cells indicate that H/ACA snoRTs (e.g. the previously identified EIF4A1 snoRT) are indeed associated with dyskerin. This resembles what occurs with other known H/ACA box snoRNAs, such as TERC and SNORA23 (notably, the independently transcribed TERC sequence is not reported as being part of any known cellular mRNA – **Figure 2B**). Then a second RIP analysis was performed after subcellular fractioning. To ensure proper fractioning and limited leakage of nuclear content in the cytoplasm we searched for both anchored (i.e. vimentin in the cytoplasm and lamin-B1 in the nucleus) and soluble (i.e. GAPDH in the cytoplasm and c-Myc in the nucleus) proteins as controls. Our results showed that the association of H/ACA snoRTs with dyskerin is particularly enriched in the cytoplasm. Worthy of note, despite its known nuclear localization, we identified detectable amounts of the 57 kDa full-length dyskerin protein in the cytoplasm (**Figure 2C**).

**Figure 2.**
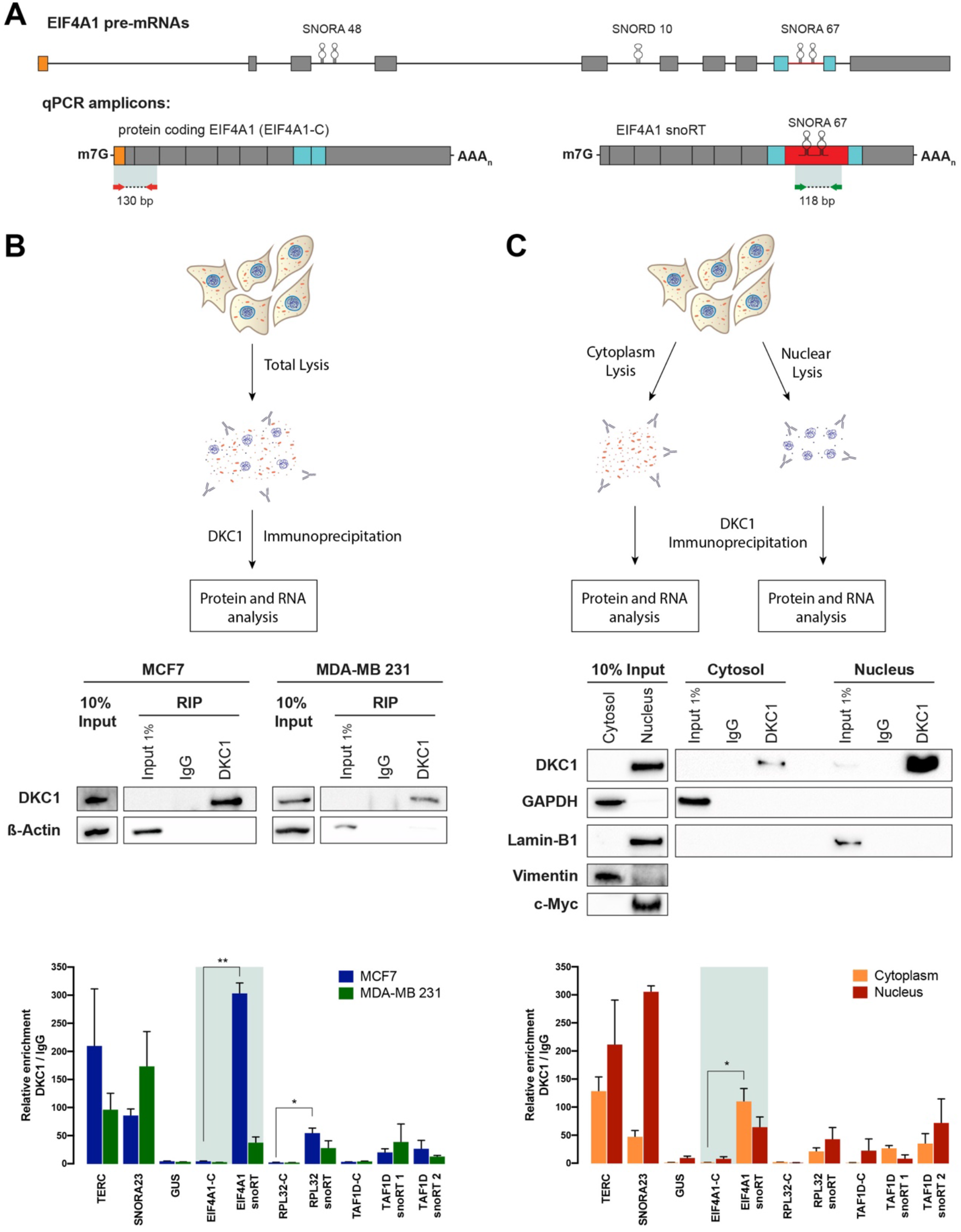
snoRT in the cytoplasm are bound by dyskerin core proteins. (A) In scale schematic overview of the EIF4A1 pre-mRNA. Introns are depicted as lines connected to exons. SNORA48, SNORD10, and SNORA67 are shown. SNORA67 intron is depicted as a red line in pre-mRNA or as a red box in the EIF4A1 snoRT intron-retaining transcript. SNORA67-flanking exons are depicted as blue boxes. The first exon of pre-mRNA is depicted as an orange box. Diagnostic qPCR amplicons are represented. The amplicon between red arrows identifies the protein coding mRNA, while the amplicon between green arrows identifies every EIF4A1 snoRTs. Primer sequences are listed in Supplementary Table S1. m7G: cap; AAAn: poly(A) tail; P: monophosphate. (B) RNA immunoprecipitation analysis of dyskerin from total cellular lysates. Top: outline of sample preparation steps. Middle: Western blot analysis of immunoprecipitated fractions from total MCF7 and MB-MDA 231 cell lysates using control IgG or anti-dyskerin antibody. A 10% input extract was used to verify the correct lysis. Bottom: RT-qPCR analysis of the known dyskerin targets (TERC, SNORA23), a known off-target (GUS), and transcripts of interest. EIF4A1 results are highlighted. Results are expressed as the fold change against immunoprecipitation with IgG. Data are shown as means ± standard error of the mean (SEM). (C) RNA immunoprecipitation analysis of dyskerin from cytoplasmic and nuclear cell lysates. Top: outline of sample preparation steps. Centre: Western blot analysis of immunoprecipitated fractions from cytoplasmic and nuclear MCF7 cell lysates using control IgG or anti-dyskerin antibody. GAPDH and Vimentin were used as cytoplasmic markers, while Lamin-B1 and c-Myc were used as nuclear markers. A 10% input extract was used to verify the correct lysis. Bottom: RT-qPCR analysis of the known dyskerin targets (TERC, SNORA23), a known off-target (GUS), and transcripts of interest. EIF4A1 results are highlighted. Results are expressed as the fold change against immunoprecipitation with IgG. Data are shown as means ± SEM. n=3 biological replicates were performed for each experiment. Paired Student’s t tests were performed on controls. *p < 0.05, **p < 0.01, ***p < 0.005, ****p < 0.001.

### H/ACA snoRTs are processed to generate cytoplasmic 5’snoRNA-ended polyadenylated transcripts bound by dyskerin

The MCF7 RNA-Seq data reported above indicate that EIF4A1 snoRT is the most regulated transcript among the identified H/ACA snoRTs after dyskerin depletion. Also, in consideration of both its relative abundancy (**Figure 3A**) and the strength of its enrichment in the dyskerin immunoprecipitation fraction (**Figure 2B**), we focused on this particular transcript to study the processing of dyskerin-bound snoRTs.

**Figure 3.**
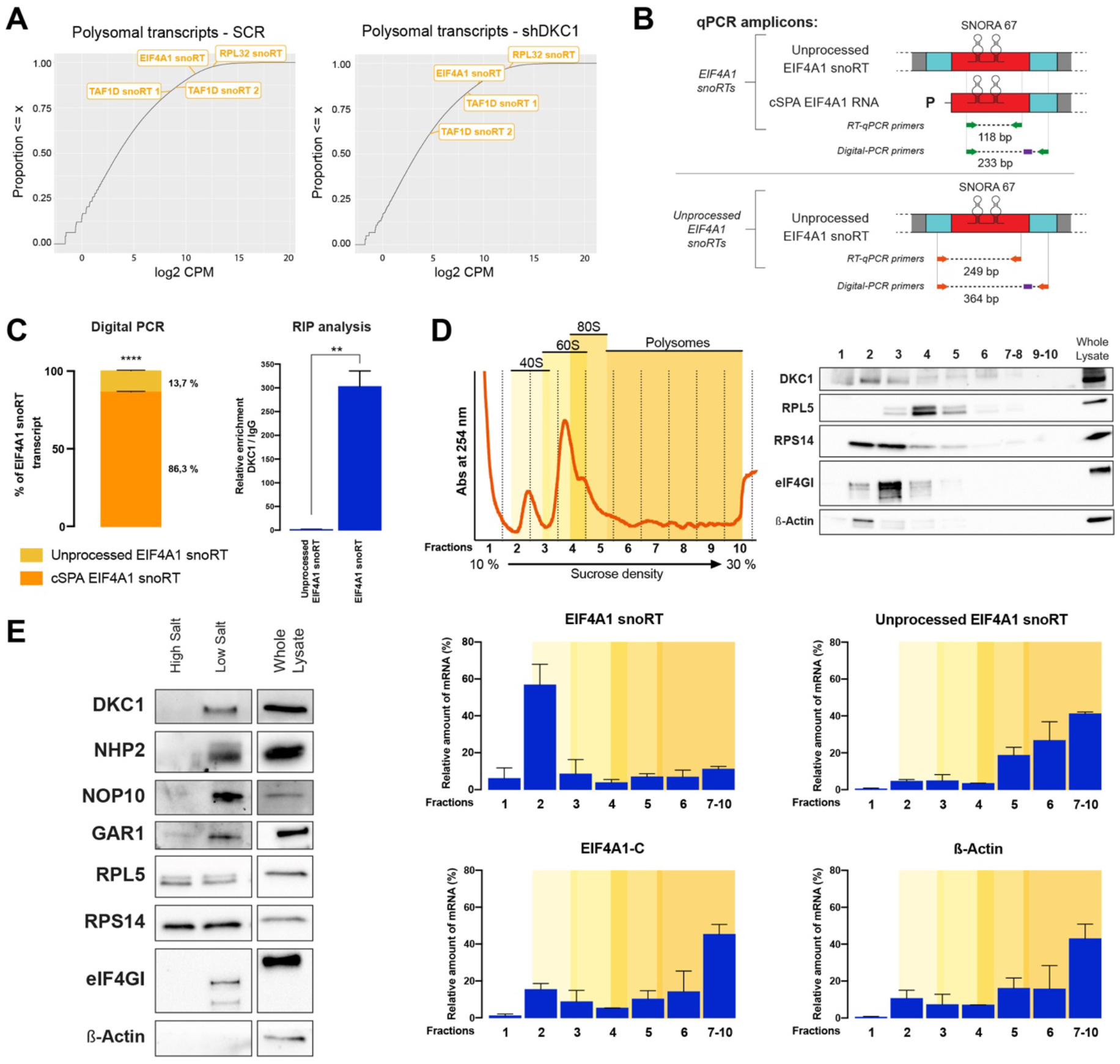
Dyskerin-bound snoRT are processed to generate 5’snoRNA-ended polyadenylated transcripts and interact with ribosomes in the cytoplasm. (A) Empirical cumulative distribution function of total cytoplasmic and polysomal-recruited transcripts. Transcripts of interest are highlighted in boxes. (B) qPCR amplicons and relative mRNA species identified by designed primers. SNORA67-flanking exons are depicted as blue boxes, while the retained intron is depicted as a red box. The amplicon between green arrows identifies every EIF4A1 snoRTs (unprocessed EIF4A1 snoRT and cSPA EIF4A1 RNA), while the amplicon between orange arrows identifies only unprocessed EIF4A1 snoRT. In purple are depicted the probes designed for digital-PCR. Primer and probe sequences are listed in Supplementary Table S1 (C) Left: Percentage representation of EIF4A1 snoRTs obtained by digital PCR absolute quantification in MCF7 cells. Data are shown as the percentage of EIF4A1 snoRT after normalization with GUS housekeeping transcript. Right: qPCR analysis of EIF4A1 snoRTs obtained by RNA immunoprecipitation of dyskerin from MCF7 total cellular lysate. The means from three biological replicates (n = 3) are shown; error bars represent SD. Paired Student’s t tests were performed on controls. *p < 0.05, **p < 0.01, ***p < 0.005, ****p < 0.001. (D) Polysome profiling analysis. Top left: representative polysome profile obtained by 10-30% sucrose density gradient centrifugation from MCF7 cells. The portions of the profile referring to different ribosomal subunits are highlighted. Top right: distribution of dyskerin and control proteins across the gradient analysed by Western blotting with specific antibodies. Bottom: distribution of transcripts of interest after RNA purification from gradient fractions obtained by RT-qPCR. Results are expressed as fraction (%) of the total amount of the transcript contained in the lysate. Data are shown as means ± SEM of two different biological replicates. (E) Ribosome purification: Western blotting analysis of purified ribosomes from MCF7 cells shows a co-purification of all pseudouridine-RNP complex (DKC1, NHP2, NOP10, GAR1). RPL5 and RPS14 are shown as a positive control for the ribosomal purification, while eIF4G is used as a control for ribosome-interacting factors.

snoRTs can be processed by non-sense mediated decay (NMD), originating <u>c</u>ytoplasmic 5’<u>s</u>noRNA ended, 3’-<u>p</u>oly<u>a</u>denylated transcripts referred to as cSPA RNA (29–31). The primers used above to identify EIF4A1 snoRT do not permit distinguishing the unprocessed full-length transcript from a possible 3’end-processed transcript. Therefore, to obtain an absolute quantification, we performed a digital RT-PCR analysis with primers capable of selectively identifying the unprocessed EIF4A1 snoRT and primers able to amplify all – unprocessed and processed – snoRTs bearing the SNORA67 containing intron (**Figure 3B**). Results on total mRNA from MCF7 cells showed that unprocessed EIF4A1 snoRT represents only 13.7% of the total EIF4A1 snoRTs. Therefore, according to the percentage difference, a processed cSPA RNA must account for the remaining 86.3% (**Figure 3C – left panel**; similar results were observed in a second cell line – **Figure S3A**). Moreover, the RIP analysis of total MCF7 cellular lysate showed that the majority of unprocessed EIF4A1 snoRT is not bound by dyskerin, while total EIF4A1 snoRT is highly enriched, indicating that dyskerin preferentially binds the processed cSPA RNA (**Figure 3C – right panel**). Accordingly, the stability of unprocessed EIF4A1 snoRT assessed after actinomycin treatment was not modified after dyskerin depletion, while that of EIF4A1 snoRT (mostly represented by the cSPA EIF4A1 RNA) was found to be strongly dyskerin-dependent (**Figure S3B)**.

### H/ACA snoRTs interact with ribosomes in the cytoplasm

To investigate how dyskerin regulates the association of H/ACA snoRTs with polysomes and their mutual interaction we performed further polysomal fractionation (**Figure 3D**). In this analysis, we characterized the association of dyskerin and H/ACA snoRTs with different cytoplasmic fractions by RT-qPCR. As previously observed, we found some of the EIF4A1 snoRT associated with polysomes. However, the majority of the EIF4A1 snoRT co-sedimented with the free 40S ribosomal subunit, as is the case with the distribution of dyskerin. Instead, the unprocessed EIF4A1 snoRT is mainly recruited to polysomal fractions, as with the canonical EIF4A1 protein-coding isoform. Therefore, these results indicate that the unprocessed EIF4A1 snoRT is translated. Furthermore, the analysis of mRNA distribution after polysomal fractionation and treatment with the translation inhibitor puromycin (**Figure S3C**) and the re-analysis of publicly available ribosome profiling datasets confirmed this finding (**Figure S3D**). On the other hand, most of the cSPA EIF4A1 RNA is associated with the small ribosomal subunit. In addition, we isolated ribosomes in low- and high-stringency conditions, demonstrating that all the RNP pseudouridylation complex core proteins are associated with the cytoplasmic ribosomes, as is the case with other ribosome-interacting factors (e.g. as with eIF4G - **Figure 3E**, similar results were observed in a second cell line – **Figure S3E**). In addition, the RIP analysis was also performed by using an antibody directed to the pseudouridylation complex core protein GAR1. Results showed that EIF4A1 snoRT is also associated with GAR1, indicating that H/ACA snoRT are bound by RNP core proteins (**Figure S3F**). All together, these results show that dyskerin binds to cSPA EIF4A1 RNA in the cytoplasm and that this complex mainly associates with the small ribosomal subunit.

### Dyskerin binds to a complex RNA interactome in the cytoplasm

To characterize the transcripts involved in the dyskerin-mediated regulation in the cytoplasm, we performed a RIP-Seq analysis of the RNAs co-immunoprecipitated by an anti-dyskerin antibody from the cytoplasmic fraction of MCF7 cells. We identified 701 significantly enriched transcripts after immunoprecipitation. The biotype distribution of the identified transcripts, according to ENSEMBL annotation (32), is shown in **Figure 4A**, left panel. Of the 701 transcripts, 115 (16.4%) are univocally classified as snoRNA and scaRNA (111 of these contain an H/ACA box). Therefore, those transcripts are expected to be bound directly by dyskerin. Of the remaining 586 transcripts, 50 (7.1%) are H/ACA snoRTs transcribed from known snoRNA host genes. The biotype distribution of the known snoRTs immunoprecipitated by dyskerin is shown in **Figures 4A** and **S4A**, right panel, where the “retained intron” biotype is the most represented (32 transcripts, 4.6%). Interestingly, these 50 H/ACA snoRTs are among the most enriched transcript isoforms after dyskerin IP (39/50 are above the median value of enrichment after IP, **Figure 4B**).

**Figure 4.**
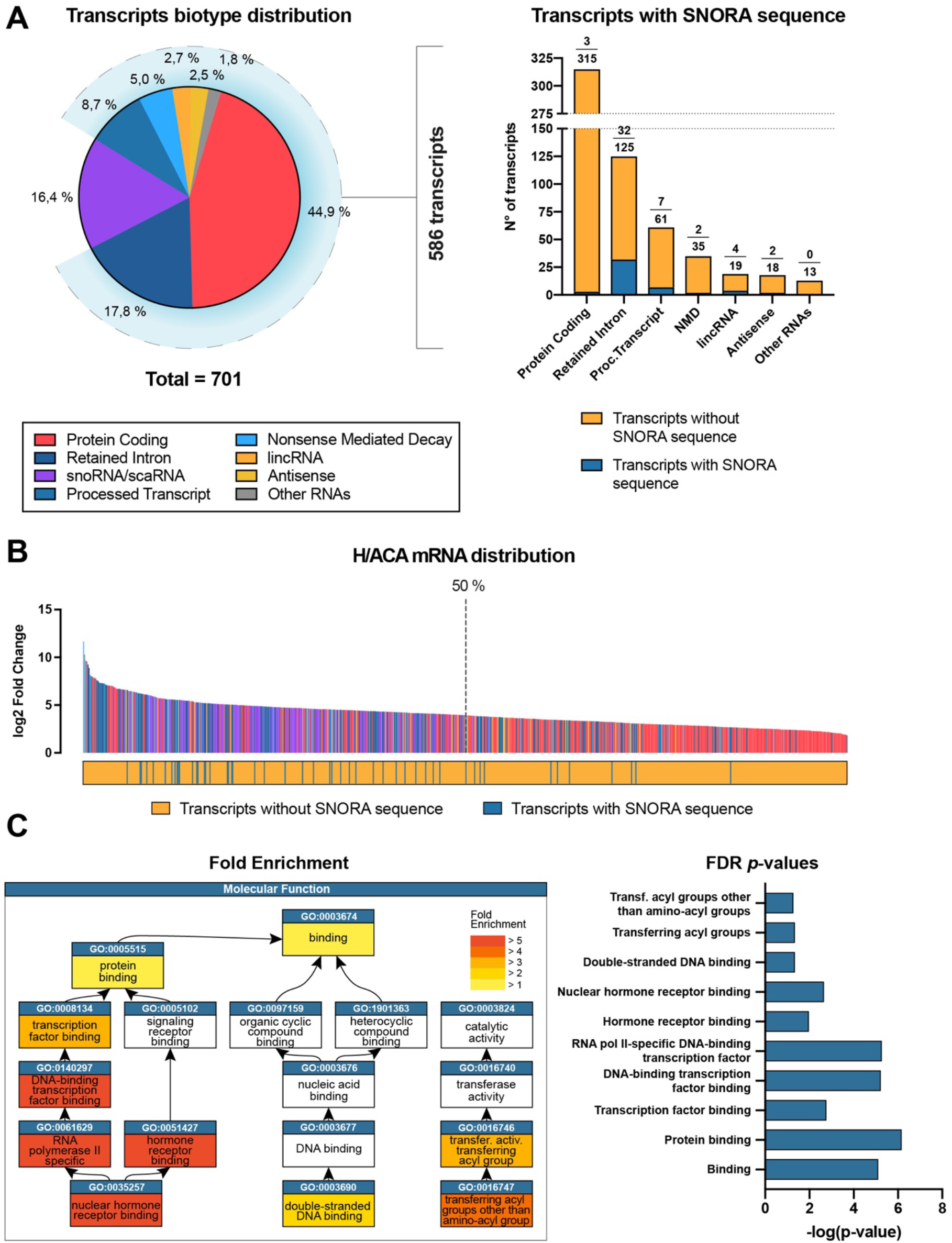
Dyskerin binds to a complex RNA interactome in the cytoplasm. (A) Left: Biotype distribution of transcripts identified by RIP-Seq analysis of RNAs co-immunoprecipitated by an anti-dyskerin antibody in a cytoplasmic MCF7 lysate. Right: Biotype distribution of H/ACA mRNAs immunoprecipitated by dyskerin from MFC7 cytoplasmic fractions. 5 independent replicates were used for RIP-seq analysis (B) Quantitative distribution of transcripts immunoprecipitated by dyskerin from MFC7 cytoplasmic fractions. The dotted vertical line is the median. The plot below the transcript distribution shows transcripts with or without SNORA sequence. (C) Fold enrichment (left) and FDR p-values (right) of the molecular function gene ontology chart of the statistically significant terms obtained from the analysis of the 238 unique protein coding transcripts immunoprecipitated by dyskerin. FDR = False Discovery Rate

However, we also found many transcripts not known to contain H/ACA snoRNA sequences to be highly enriched by RIP-seq. To investigate the effects of dyskerin modulation on these transcripts, we once again looked at our RNA-Seq data, focusing only on the 701 dyskerin-associated transcripts. Over all, these transcripts appear to be significantly regulated after dyskerin depletion compared to the rest of the transcriptome (**Figure S4B**), suggesting that they may be regulated by this protein. To obtain functional insights into this regulation, we performed a Gene Ontology analysis on the list of genes that transcribe for the mRNAs that we found associated with dyskerin. To avoid the possible confounding effect deriving from the cases where the association with dyskerin occurs through non-coding transcript isoforms (e.g. *EIF4A1*), we focused only on 238 genes corresponding to the 312 immunoprecipitated protein-coding transcripts. The most strongly enriched molecular function turned out to be “nuclear hormone receptor binding” (**Figure 4C, Figure S4C**).

### Dyskerin modulates nuclear hormone receptor-mediated dependence in human breast cancer cells

To characterize the role of the above-identified dyskerin interactions, we investigated the effect of removing the ligands of nuclear hormone receptors from the serum used in *in vitro* experiments by charcoal treatment in shDKC1 MCF7 cells and their relevant controls. The results obtained indicate that when cultured with the addition of full serum, dyskerin depletion did not affect the expression of selected known direct targets of the two main hormone receptors active in MCF7 cells, oestrogen receptor – ER – and progesterone receptor – PGR. Conversely, under charcoal treatment conditions, most of the tested targets were influenced by dyskerin depletion (**Figure 5A**). These results offer a preliminary indication that in breast cancer cells the lack of dyskerin-mediated regulation may confer some degree of oestrogen and progesterone independence. To evaluate this possibility, we retrospectively analysed a series of primary breast carcinomas available at our institution. In this series, to obtain indications on the interaction of dyskerin with the cytoplasmic mRNAs identified in the RIP-Seq analysis, we used RT-qPCR to quantify the levels of the EIF4A1 snoRT, which is one of the most abundant and quantitatively regulated dyskerin-bound cytoplasmic RNAs. In this analysis, therefore, EIF4A1 snoRT levels were considered to be a proxy for dyskerin cytoplasmic RNA-binding. In the analysed series, EIF4A1 snoRT levels were significantly correlated with DKC1 mRNA levels (**Figure S5A**). In addition, the results obtained showed that EIF4A1 snoRT levels were significantly directly related to the ER and PGR labelling index obtained at the time of diagnosis by immunohistochemistry (**Figure 5B**). EIF4A1 snoRT levels in tumour specimens also proved to be strongly associated with patients’ specific survival (**Figure 5C**), as is the case with the ER and PGR status (33). Importantly, the prognostic value of EIF4A1 snoRT was particularly evident in ER-positive and PGR-positive cases (**Figure 5C**), again suggesting that the alteration in dyskerin cytoplasmic functions may be associated with oestrogen and progesterone independence. To obtain experimental *in vivo* evidence on this point, we xenografted nude mice subcutaneously with MCF7 shDKC1 and control cells in the absence of any additional oestrogen treatment. In fact, the parental MCF7 cell line is known to be highly oestrogen-dependent and to be tumorigenic in nude mice subcutaneously when they are continuously treated with oestrogen supplementation after the xenografting procedure (34). Control cells in these conditions consistently generated tumours only in a minimal number of cases and after a long observation time (3/10 after 26 weeks). In contrast, shDKC1 cells displayed a significantly higher tumorigenic potential (6/10, p= 0.04) (**Figure 5D**). All together, these results indicate that the dysregulation of the cytoplasmic RNA-binding by dyskerin alters the dependence of breast cancer cells on nuclear hormone receptor ligands.

**Figure 5.**
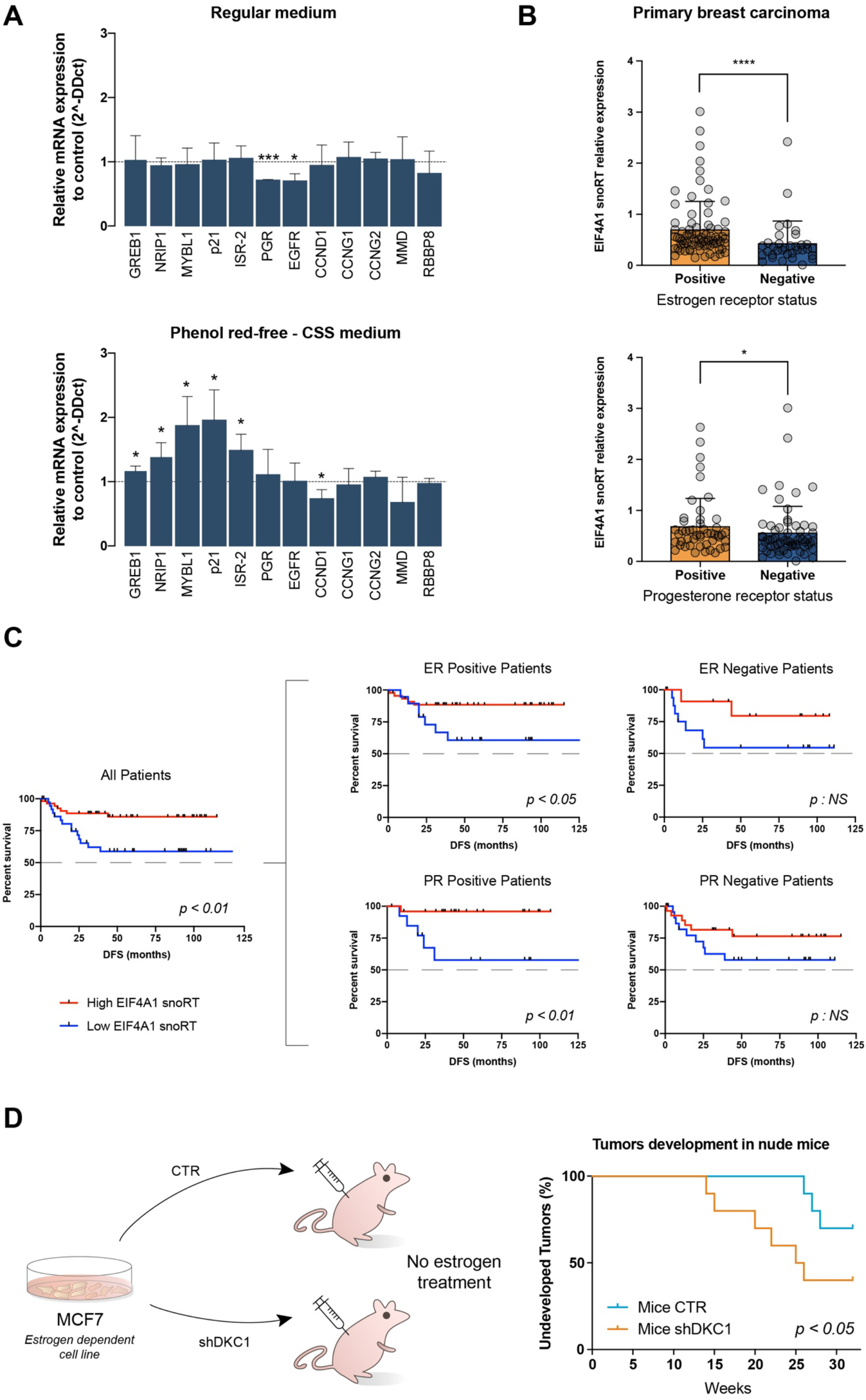
Dyskerin levels modify nuclear hormone receptor-mediated dependence in human breast cancer cells. (A) qPCR analysis of selected known target transcripts of oestrogen and progesterone receptors, observed under standard medium condition or with a phenol red-free and charcoal-stripped serum (CSS) medium condition. Data are shown as a fold change of shDKC1 MCF7 cells relative to their controls. The means from three biological replicates (n = 3) are displayed, error bars represent SD. Paired Student’s t tests were performed on controls. (B) EIF4A1 snoRT relative expression obtained by RT-qPCR of RNA extracted from 111 breast cancer tissues. Patients are divided into positive and negative for oestrogen receptors (top) or progesterone receptors (bottom). Error bars represent SD. Mann-Whitney tests were performed on controls. (C) Kaplan-Meier survival curves for breast cancer patients. Patients are divided between high and low EIF4A1 snoRT expression (separated by the median value EIF4A1 snoRT expression) obtained by qPCR analysis. The left plot shows the curves of all patients (n = 90) while the right plots show only the curves for oestrogen receptor-positive (n = 62) and -negative (n = 27) (top), or progesterone receptor-positive (n = 38) and -negative (n = 51) (bottom) patients. (D) Experimental plan (left). MCF7 cells (which are dependent on oestrogen for their tumorigenic potential) were injected subcutaneously into mice (10 replicates for CTR and 10 replicates for shDKC1). No supplementary oestrogen was administered. Kaplan-Meier curves (right) show the development of tumours in nude mice xenografted with control and shDKC1 MCF7 cells. *p < 0.05, **p < 0.01, ***p < 0.005, ****p < 0.001, NS (not significant).

## DISCUSSION

The present study was initially aimed at identifying genes whose expression is modulated by dyskerin. The unbiased analysis performed on total and polysomal RNA fractions identified a group of transcripts retaining introns with H/ACA box snoRNA sequences, that we defined as snoRTs, impacted by dyskerin depletion. However, known IRES-containing dyskerin translational targets such as Bcl-XL, XIAP, p27 (14), P53 (16), and VEGF (17) were not identified in our analysis. This might be explained by the fact that in order to identify undescribed translational targets in our study, exponentially growing cells were analysed without inducing the stress conditions that may elicit cap-independent translation. H/ACA snoRTs were identified in the cytoplasm of cells (and, to some extent, on polysomes), and turned out to be bound by dyskerin. At this regard it is worth noting that, although in the fractioning conditions we employed a minimal leakage of soluble nuclear components in the cytoplasm cannot be completely excluded, our fractioning controls indicate that this occurrence is actually extremely limited. Although dyskerin is well-known to be localized in the nucleus, a previous study identified a cytoplasmic isoform of this protein characterized by a lower molecular weight (35). In our analyses, however, the molecular weight of the dyskerin product identified in the cytoplasm corresponds to that of the canonical full-length protein. The association of the predominantly nuclear isoform of dyskerin with polysomes and ribosomal subunits strongly support that its presence in the cytoplasm is not due to a simple leakage during the fractioning procedure.

This indicates that, in addition to its nuclear localization, a fraction of cellular dyskerin is involved in mRNA binding in the cytoplasm. Our observations match what has been described for the 2’-O-methyltransferase core proteins, which are found bound to a retained intron containing a C/D snoRNA sequence in a snoRT transcribed from the NOP56 gene (31). It has also been reported that this specific transcript is processed by NMD to a cSPA RNA. Our results obtained on the EIF4A1 snoRT suggest that this may also occur for H/ACA snoRT in association with pseudouridylation core proteins. In fact, the EIF4A1 snoRT was found to be recruited to polysomes, a step necessary for NMD. A significant fraction of it, however, was detected in association with the free 40S subunit. In particular, the comparison of the recruitment of the different transcript isoforms identified by qPCR indicates that the large majority of the EIF4A1 snoRT associated with 40S consists of the processed cSPA EIF4A1 RNA. In this regard, it is worth noting that the canonical SNORA67 is known to target the sequences flanking U1445 on 18S rRNA, which are relatively exposed on the outer surface of the 40S ribosomal subunit (**Figure S6**) by complementary base pairing (36). This kind of interaction might also be involved in the association between EIF4A1 snoRT and the 40S subunit.

On the basis of these previously uncovered dyskerin functions in the cytoplasm, we then comprehensively characterized the RNA molecules interacting with dyskerin in the cytoplasm via a RIP-seq analysis from cellular cytoplasmic fractions. We identified 701 significantly enriched transcripts, among which 50 H/ACA snoRTs were found. With very few exceptions, intron retention introduces a premature stop codon into the host gene transcript, thus preventing protein expression. However, 312 protein-coding transcripts not containing any known H/ACA box snoRNA sequence were also co-immunoprecipitated. The nature of the interaction of these mRNAs with dyskerin remains to be characterized. In particular, it would be interesting to know whether they interact directly (possibly through the recognition of yet unrecognized binding domains), or if the contact between them is mediated by other transcripts bound by dyskerin, such as those retaining H/ACA-containing introns. Independently of the underlying type of interaction, a significant fraction of immunoprecipitated transcripts encode for proteins involved in mediating the effect of nuclear hormone receptor ligands. Interestingly, our results suggest that in breast cancer cells, the lack of cytoplasmic dyskerin functions may provide a way for luminal hormone-receptor positive tumours to escape the hormone dependence, thus becoming more aggressive. Such an effect may contribute to explaining the well-known role of dyskerin in mammary tumorigenesis (2,11,17). In this regard, however, we recognize that it is not always possible, when looking at previous studies, to dissect dyskerin cytoplasmic-specific functions from the well-established effects on telomerase activity and rRNA pseudouridylation.

Our results may be of importance also for X-linked dyskeratosis congenita, caused by point mutations of the DKC1 gene (1). In the present study, we do not investigate the effects of these mutations on the described cytoplasmic functions of dyskerin. It is worth remembering, however, that one important therapeutic option for patients with dyskeratosis congenita, as well in all the so-called telomeropathies, is treatment with androgen derivatives (37). Therefore, the investigation of androgen response, specifically in patients bearing DKC1 mutations in future studies, appears useful. In addition, a potential nuclear hormone receptor independence in patients with DKC1 mutations may also play a role in their well-known susceptibility to developing malignancies (38). The effect of pathogenic DKC1 mutations on this aspect also requires dedicated investigations.

In conclusion, our findings characterize for the first time a dyskerin-dependent mRNA post-transcriptional regulation mechanism occurring in the cytoplasm which may be significant for nuclear hormone receptor function and affect the behaviour of breast cancer cells.

## DATA AVAILABILITY

The RNA-seq and RIP-seq datasets were deposited in the Gene Expression Omnibus with ID GSE161481.

## SUPPLEMENTARY DATA

Supplementary Data are available separately

## ACKNOWLEDGEMENTS

The Authors are grateful to Dario Rizzotto for technical advice and to the Bassot Family for their support.

## REFERENCES

1. Heiss, N.S., Knight, S.W., Vulliamy, T.J., Klauck, S.M., Wiemann, S., Mason, P.J., Poustka, A. and Dokal, I. (1998) X-linked dyskeratosis congenita is caused by mutations in a highly conserved gene with putative nucleolar functions. Nat Genet, 19, 32–38.

2. Montanaro, L., Brigotti, M., Clohessy, J., Barbieri, S., Ceccarelli, C., Santini, D., Taffurelli, M., Calienni, M., Teruya-Feldstein, J., Trerè, D. et al. (2006) Dyskerin expression influences the level of ribosomal RNA pseudo-uridylation and telomerase RNA component in human breast cancer. J Pathol, 210, 10–18.

3. Mitchell, J.R., Wood, E. and Collins, K. (1999) A telomerase component is defective in the human disease dyskeratosis congenita. Nature, 402, 551–555.

4. Meier, U.T. and Blobel, G. (1994) NAP57, a mammalian nucleolar protein with a putative homolog in yeast and bacteria. J Cell Biol, 127, 1505–1514.

5. Bousquet-Antonelli, C., Henry, Y., G’Elugne J, P., Caizergues-Ferrer, M. and Kiss, T. (1997) A small nucleolar RNP protein is required for pseudouridylation of eukaryotic ribosomal RNAs. Embo j, 16, 4770–4776.

6. Henras, A., Henry, Y., Bousquet-Antonelli, C., Noaillac-Depeyre, J., Gélugne, J.P. and Caizergues-Ferrer, M. (1998) Nhp2p and Nop10p are essential for the function of H/ACA snoRNPs. Embo j, 17, 7078–7090.

7. Kiss, A.M., Jády, B.E., Bertrand, E. and Kiss, T. (2004) Human box H/ACA pseudouridylation guide RNA machinery. Mol Cell Biol, 24, 5797–5807.

8. Darzacq, X., Kittur, N., Roy, S., Shav-Tal, Y., Singer, R.H. and Meier, U.T. (2006) Stepwise RNP assembly at the site of H/ACA RNA transcription in human cells. J Cell Biol, 173, 207–218.

9. Richard, P. and Kiss, T. (2006) Integrating snoRNP assembly with mRNA biogenesis. EMBO Rep, 7, 590–592.

10. Tollervey, D. and Kiss, T. (1997) Function and synthesis of small nucleolar RNAs. Curr Opin Cell Biol, 9, 337–342.

11. Ruggero, D., Grisendi, S., Piazza, F., Rego, E., Mari, F., Rao, P.H., Cordon-Cardo, C. and Pandolfi, P.P. (2003) Dyskeratosis congenita and cancer in mice deficient in ribosomal RNA modification. Science, 299, 259–262.

12. Bellodi, C., McMahon, M., Contreras, A., Juliano, D., Kopmar, N., Nakamura, T., Maltby, D., Burlingame, A., Savage, S.A., Shimamura, A. et al. (2013) H/ACA small RNA dysfunctions in disease reveal key roles for noncoding RNA modifications in hematopoietic stem cell differentiation. Cell Rep, 3, 1493–1502.

13. Taoka, M., Nobe, Y., Yamaki, Y., Sato, K., Ishikawa, H., Izumikawa, K., Yamauchi, Y., Hirota, K., Nakayama, H., Takahashi, N. et al. (2018) Landscape of the complete RNA chemical modifications in the human 80S ribosome. Nucleic Acids Res, 46, 9289–9298.

14. Yoon, A., Peng, G., Brandenburger, Y., Zollo, O., Xu, W., Rego, E. and Ruggero, D. (2006) Impaired control of IRES-mediated translation in X-linked dyskeratosis congenita. Science, 312, 902–906.

15. Bellodi, C., Kopmar, N. and Ruggero, D. (2010) Deregulation of oncogene-induced senescence and p53 translational control in X-linked dyskeratosis congenita. Embo j, 29, 1865–1876.

16. Montanaro, L., Calienni, M., Bertoni, S., Rocchi, L., Sansone, P., Storci, G., Santini, D., Ceccarelli, C., Taffurelli, M., Carnicelli, D. et al. (2010) Novel dyskerin-mediated mechanism of p53 inactivation through defective mRNA translation. Cancer Res, 70, 4767–4777.

17. Rocchi, L., Pacilli, A., Sethi, R., Penzo, M., Schneider, R.J., Treré, D., Brigotti, M. and Montanaro, L. (2013) Dyskerin depletion increases VEGF mRNA internal ribosome entry site-mediated translation. Nucleic Acids Res, 41, 8308–8318.

18. Penzo, M., Rocchi, L., Brugiere, S., Carnicelli, D., Onofrillo, C., Couté, Y., Brigotti, M. and Montanaro, L. (2015) Human ribosomes from cells with reduced dyskerin levels are intrinsically altered in translation. Faseb j, 29, 3472–3482.

19. Guerrieri, A.N., Zacchini, F., Onofrillo, C., Di Viggiano, S., Penzo, M., Ansuini, A., Gandin, I., Nobe, Y., Taoka, M., Isobe, T. et al. (2020) DKC1 Overexpression Induces a More Aggressive Cellular Behavior and Increases Intrinsic Ribosomal Activity in Immortalized Mammary Gland Cells. Cancers, 12.

20. Penzo, M., Carnicelli, D., Montanaro, L. and Brigotti, M. (2016) A reconstituted cell-free assay for the evaluation of the intrinsic activity of purified human ribosomes. Nat Protoc, 11, 1309–1325.

21. Wu, P., Wang, X., Li, F., Qi, B., Zhu, H., Liu, S., Cui, Y. and Chen, J. (2008) Growth suppression of MCF-7 cancer cell-derived xenografts in nude mice by caveolin-1. Biochem Biophys Res Commun, 376, 215–220.

22. Bolger, A.M., Lohse, M. and Usadel, B. (2014) Trimmomatic: a flexible trimmer for Illumina sequence data. Bioinformatics, 30, 2114–2120.

23. Dobin, A., Davis, C.A., Schlesinger, F., Drenkow, J., Zaleski, C., Jha, S., Batut, P., Chaisson, M. and Gingeras, T.R. (2013) STAR: ultrafast universal RNA-seq aligner. Bioinformatics, 29, 15–21.

24. Patro, R., Duggal, G., Love, M.I., Irizarry, R.A. and Kingsford, C. (2017) Salmon provides fast and bias-aware quantification of transcript expression. Nat Methods, 14, 417–419.

25. Love, M.I., Huber, W. and Anders, S. (2014) Moderated estimation of fold change and dispersion for RNA-seq data with DESeq2. Genome Biol, 15, 550.

26. Anders, S., Reyes, A. and Huber, W. (2012) Detecting differential usage of exons from RNA-seq data. Genome Res, 22, 2008–2017.

27. Alexa, A. and Rahnenfuhrer, J. (2020).

28. Chenchik, A. (1998) Generation and use of high-quality cDNA from small amounts of total RNA by SMART PCR. Gene cloning and analysis by RT-PCR.

29. Lykke-Andersen, S., Chen, Y., Ardal, B.R., Lilje, B., Waage, J., Sandelin, A. and Jensen, T.H. (2014) Human nonsense-mediated RNA decay initiates widely by endonucleolysis and targets snoRNA host genes. Genes Dev, 28, 2498–2517.

30. Wu, H., Yin, Q.F., Luo, Z., Yao, R.W., Zheng, C.C., Zhang, J., Xiang, J.F., Yang, L. and Chen, L.L. (2016) Unusual Processing Generates SPA LncRNAs that Sequester Multiple RNA Binding Proteins. Mol Cell, 64, 534–548.

31. Lykke-Andersen, S., Ardal, B.K., Hollensen, A.K., Damgaard, C.K. and Jensen, T.H. (2018) Box C/D snoRNP Autoregulation by a cis-Acting snoRNA in the NOP56 Pre-mRNA. Mol Cell, 72, 99–111.e115.

32. Zerbino, D.R., Achuthan, P., Akanni, W., Amode, M.R., Barrell, D., Bhai, J., Billis, K., Cummins, C., Gall, A., Girón, C.G. et al. (2018) Ensembl 2018. Nucleic Acids Res, 46, D754–d761.

33. Hawkins, R.A., Tesdale, A.L., Killen, M.E., Jack, W.J., Chetty, U., Dixon, J.M., Hulme, M.J., Prescott, R.J., McIntyre, M.A. and Miller, W.R. (1996) Prospective evaluation of prognostic factors in operable breast cancer. Br J Cancer, 74, 1469–1478.

34. Levy, J.A., White, A.C. and McGrath, C.M. (1982) Growth and histology of a human mammary-carcinoma cell line at different sites in the athymic mouse. Br J Cancer, 45, 375–383.

35. Angrisani, A., Turano, M., Paparo, L., Di Mauro, C. and Furia, M. (2011) A new human dyskerin isoform with cytoplasmic localization. Biochim Biophys Acta, 1810, 1361–1368.

36. Ganot, P., Bortolin, M.L. and Kiss, T. (1997) Site-specific pseudouridine formation in preribosomal RNA is guided by small nucleolar RNAs. Cell, 89, 799–809.

37. Calado, R.T. and Clé, D.V. (2017) Treatment of inherited bone marrow failure syndromes beyond transplantation. Hematology Am Soc Hematol Educ Program, 2017, 96–101.

38. Alter, B.P., Giri, N., Savage, S.A. and Rosenberg, P.S. (2018) Cancer in the National Cancer Institute inherited bone marrow failure syndrome cohort after fifteen years of follow-up. Haematologica, 103, 30–39.

