## Supplementary Information for "The pseudouridine synthase dyskerin binds to cytoplasmic H/ACA-box snoRNA retaining transcripts affecting nuclear hormone receptor dependence"

**A**

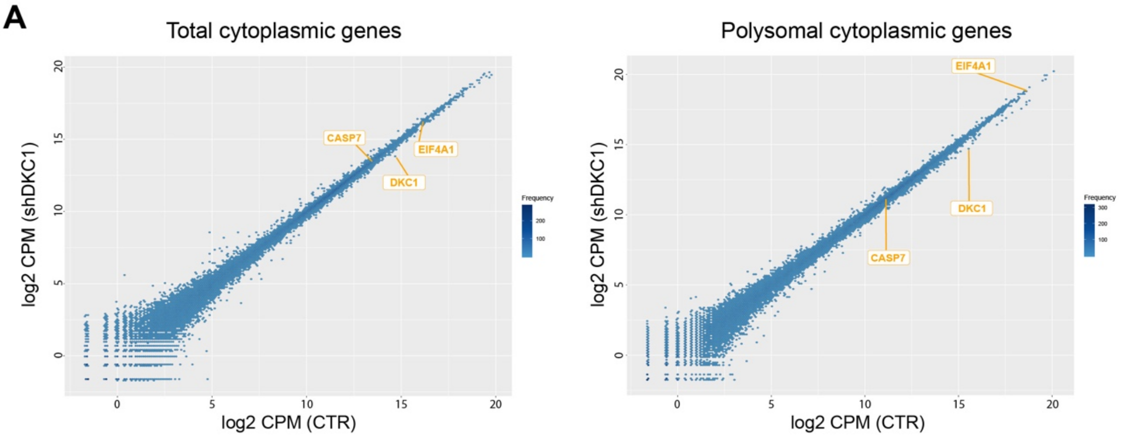

| Upregulated Genes | | |
|---|---|---|
| Gene | Log2 FC | pValue |

| Downregulated Genes | | |
|---|---|---|
| Gene | Log2 FC | pValue |
| DKC1 | -0.755 | 2.76E-05 |

| Upregulated Genes | | |
|---|---|---|
| Gene | Log2 FC | pValue |

| Downregulated Genes | | |
|---|---|---|
| Gene | Log2 FC | pValue |
| DKC1 | -0.732 | 8.71E-05 |

**B**

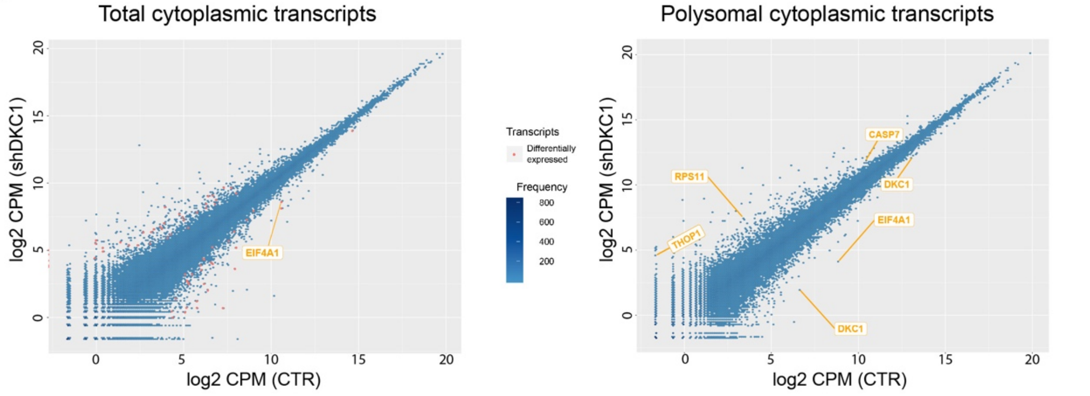

**DKC1 silenced total mRNA vs control total mRNA - Transcript Analysis**

| Upregulated Genes | | |
|---|---|---|
| Gene | Log2 FC | pValue |
| ZNF584 | 1.72 | 0.00012 |
| TYK2 | 2.55 | 0.00018 |
| ING4 | 2.16 | 0.00058 |
| HHEX | 2.21 | 0.00122 |
| HAUS7 | 2.32 | 0.00223 |
| PSMB10 | 2.35 | 0.00334 |
| HLA-B | 2.34 | 0.00334 |
| CRKL | 2.28 | 0.00641 |
| ARF5 | 2.18 | 0.00734 |
| C11orf74 | 2.10 | 0.00734 |
| CD59 | 2.21 | 0.00779 |
| ETV7 | 2.07 | 0.00779 |
| USP4 | 2.06 | 0.00973 |
| CAST | 1.73 | 0.01330 |
| BSG | 2.07 | 0.01349 |
| THBS3 | 1.60 | 0.01349 |
| CDK20 | 1.73 | 0.01409 |
| ADAM15 | 1.44 | 0.01409 |
| ATP5O | 1.82 | 0.01864 |
| ANKZF1 | 2.09 | 0.01893 |
| TRAPPC9 | 2.08 | 0.01893 |
| MKLN1-AS | 2.07 | 0.01894 |
| SPAG4 | 1.53 | 0.02029 |
| HMGCL | 1.65 | 0.02157 |
| MAP2K5 | 1.98 | 0.02267 |
| PCYT2 | 2.04 | 0.02270 |
| STX1A | 1.57 | 0.02270 |
| DERA | 1.83 | 0.03586 |
| RAC3 | 1.96 | 0.03951 |
| HERPUD1 | 1.93 | 0.03951 |
| ATP1B3 | 1.80 | 0.03951 |
| FDFT1 | 1.53 | 0.04067 |
| CAST | 1.82 | 0.04726 |
| AES | 1.74 | 0.04726 |
| RPS11 | 1.25 | 0.04836 |
| AC016394.1 | 0.00 | 0.04860 |
| HEXB | 1.73 | 0.04889 |
| KHSRP | 1.10 | 0.04889 |

| Downregulated Genes | | |
|---|---|---|
| Gene | Log2 FC | pValue |
| DKC1 | -0.76 | 0.00048 |
| RBM28 | -1.99 | 0.00058 |
| RNF114 | -2.28 | 0.00061 |
| SMARCA2 | -2.39 | 0.00223 |
| RAB7A | -2.25 | 0.00349 |
| EIF4A1 | -1.69 | 0.00734 |
| FBXO24 | -1.84 | 0.00734 |
| REXO2 | -1.91 | 0.00734 |
| TMCC1-AS1 | -2.17 | 0.00734 |
| SBDS | -2.24 | 0.00734 |
| TPRA1 | -1.58 | 0.00779 |
| RAE1 | -1.81 | 0.01572 |
| WAS | -1.94 | 0.01870 |
| PLXNB2 | -1.01 | 0.01893 |
| NAXE | -1.86 | 0.02714 |
| MVK | -1.21 | 0.02987 |
| CCDC47 | -1.97 | 0.03013 |
| P2RX4 | -1.85 | 0.03335 |
| CARD14 | -1.47 | 0.03530 |
| DKC1 | -1.53 | 0.03716 |
| NUMB | -1.23 | 0.03951 |
| SON | -1.96 | 0.03951 |
| NT5DC2 | -1.49 | 0.04067 |
| DPAGT1 | -1.69 | 0.04195 |
| EIF3J | -1.94 | 0.04513 |
| SLC6A14 | -1.63 | 0.04726 |
| NSUN5P1 | -1.76 | 0.04726 |
| ASNS | -1.78 | 0.04726 |
| LMAN2 | -1.91 | 0.04742 |
| E2F4 | -1.92 | 0.04860 |
| MAP3K4 | -1.89 | 0.04889 |

**DKC1 silenced polysomal mRNA vs control polysomal mRNA - Transcript Analysis**

| Upregulated Genes | | |
|---|---|---|
| Gene | Log2 FC | pValue |
| CASP7 | 1.24 | 0.00063 |
| RPS11 | 1.59 | 0.00700 |
| THOP1 | 2.27 | 0.00975 |

| Downregulated Genes | | |
|---|---|---|
| Gene | Log2 FC | pValue |
| EIF4A1 | -3.55 | 0.00049 |
| DKC1 | -0.73 | 0.00264 |
| DKC1 | -2.01 | 0.00278 |

**C**

### Total cytoplasmic transcripts (exon analysis)

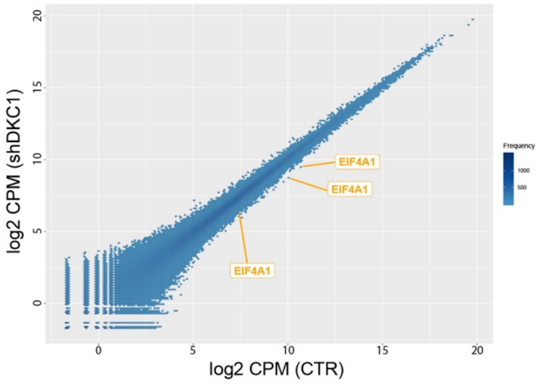

**DKC1 silenced total mRNA vs control total mRNA - Exon Analysis**

**Upregulated Genes**

| Gene | Log2 FC | pValue |
|---|---|---|
| STOML1 | 1.57 | 0.016923 |
| TEX22+MTA1 | 1.24 | 0.008611 |
| CUL3 | 1.05 | 0.001242 |
| PAM | 1.02 | 0.006946 |
| ANKZF1 | 1.00 | 0.040349 |
| RPL32+SNORA7A | 0.98 | 0.000298 |
| PAM | 0.86 | 0.030437 |
| HELLS+AL138759.1 | 0.82 | 0.00398 |
| AC134407.3+BPTF | 0.81 | 0.001464 |
| HSPH1 | 0.70 | 0.039873 |
| IQGAP1 | 0.70 | 0.047061 |
| FAT1 | 0.67 | 0.029353 |
| ARHGAP21 | 0.60 | 0.016716 |
| C8orf88 | 0.57 | 0.031289 |
| PRNP | 0.53 | 0.000574 |
| PSMA4 | 0.49 | 0.012853 |
| FGD6 | 0.44 | 0.002605 |
| BCLAF1 | 0.37 | 3.95E-05 |
| ESRP1 | 0.36 | 1.4E-05 |
| ARL6IP1+AC138811.2+RPS15A | 0.29 | 6.96E-07 |
| FAF2 | 0.28 | 0.047061 |
| ITPRIP | 0.20 | 0.0006 |

**Downregulated Genes**

| Gene | Log2 FC | pValue |
|---|---|---|
| CKS1B | -0.24 | 0.039873 |
| PAPOLA | -0.28 | 0.037781 |
| LSM14B | -0.30 | 0.047061 |
| SNORD38B+RPS8+SNORD38A+SNORD55+SNORD38B | -0.44 | 0.0443 |
| SUPT20H | -0.52 | 0.040045 |
| TGFBRAP1 | -0.72 | 0.0227 |
| SNORD10+SNORA48+CD68+SENP3+AC016876.2+SENP3-EIF4A1+SNORA67+EIF4A1 | -0.76 | 7.68E-11 |
| STAG3L5P+PILRA+PILRB+MIR6840+PVRIG2P+STAG3L5P-PVRIG2P-PILRB | -1.01 | 0.047295 |
| F11R+AL591806.3+TSTD1 | -1.02 | 0.00301 |
| GLG1 | -1.05 | 0.003947 |
| THOC7 | -1.09 | 0.0227 |
| MYO5C | -1.11 | 0.028883 |
| HNRNPD | -1.13 | 0.013817 |
| ABCB8 | -1.30 | 0.038844 |
| SNORD10+SNORA48+CD68+SENP3+AC016876.2+SENP3-EIF4A1+SNORA67+EIF4A1 | -1.31 | 0.000137 |
| SNORD10+SNORA48+CD68+SENP3+AC016876.2+SENP3-EIF4A1+SNORA67+EIF4A1 | -1.32 | 0.012727 |
| SNORD10+SNORA48+CD68+SENP3+AC016876.2+SENP3-EIF4A1+SNORA67+EIF4A1 | -1.34 | 3.57E-06 |
| MIR3175+LINC01578+CHD2+AC013394.1 | -1.40 | 0.032635 |
| NSD2 | -1.45 | 0.038844 |
| HAGHL | -2.13 | 0.001242 |

### Polysomal cytoplasmic transcripts (exon analysis)

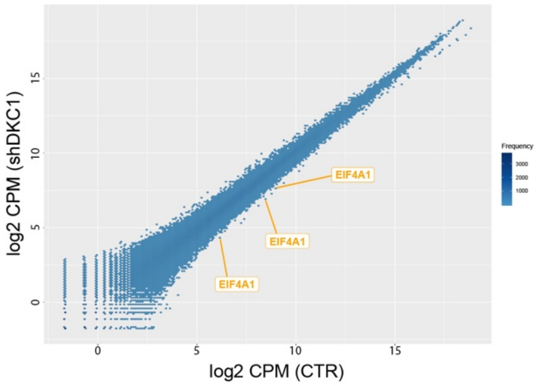

**DKC1 silenced polysomal mRNA vs control polysomal mRNA - Exon Analysis**

**Upregulated Genes**

| Gene | Log2 FC | pValue |
|---|---|---|
| RPL32+SNORA7A | 1.16 | 0.00052 |
| ZBTB43 | 0.24 | 0.00008 |

**Downregulated Genes**

| Gene | Log2 FC | pValue |
|---|---|---|
| FAM120A | -0.19 | 0.0120758 |
| TAF1D+MIR1304+SNORA25+SNORA8+SNORA18+SNORA1+SNORA32+SNORA40+SNORD5 | -0.98 | 0.0178653 |
| DYNC1H1 | -0.71 | 0.0261334 |
| SNORA56+MIR664B+DKC1+SNORA36A | -0.83 | 0.0272186 |
| SNORD10+SNORA48+CD68+SENP3+AC016876.2+SENP3-EIF4A1+SNORA67+EIF4A1 | -1.88 | 0.0261341 |
| SNORD10+SNORA48+CD68+SENP3+AC016876.2+SENP3-EIF4A1+SNORA67+EIF4A1 | -2.18 | 0.0120758 |
| SNORD10+SNORA48+CD68+SENP3+AC016876.2+SENP3-EIF4A1+SNORA67+EIF4A1 | -2.09 | 0.0000047 |

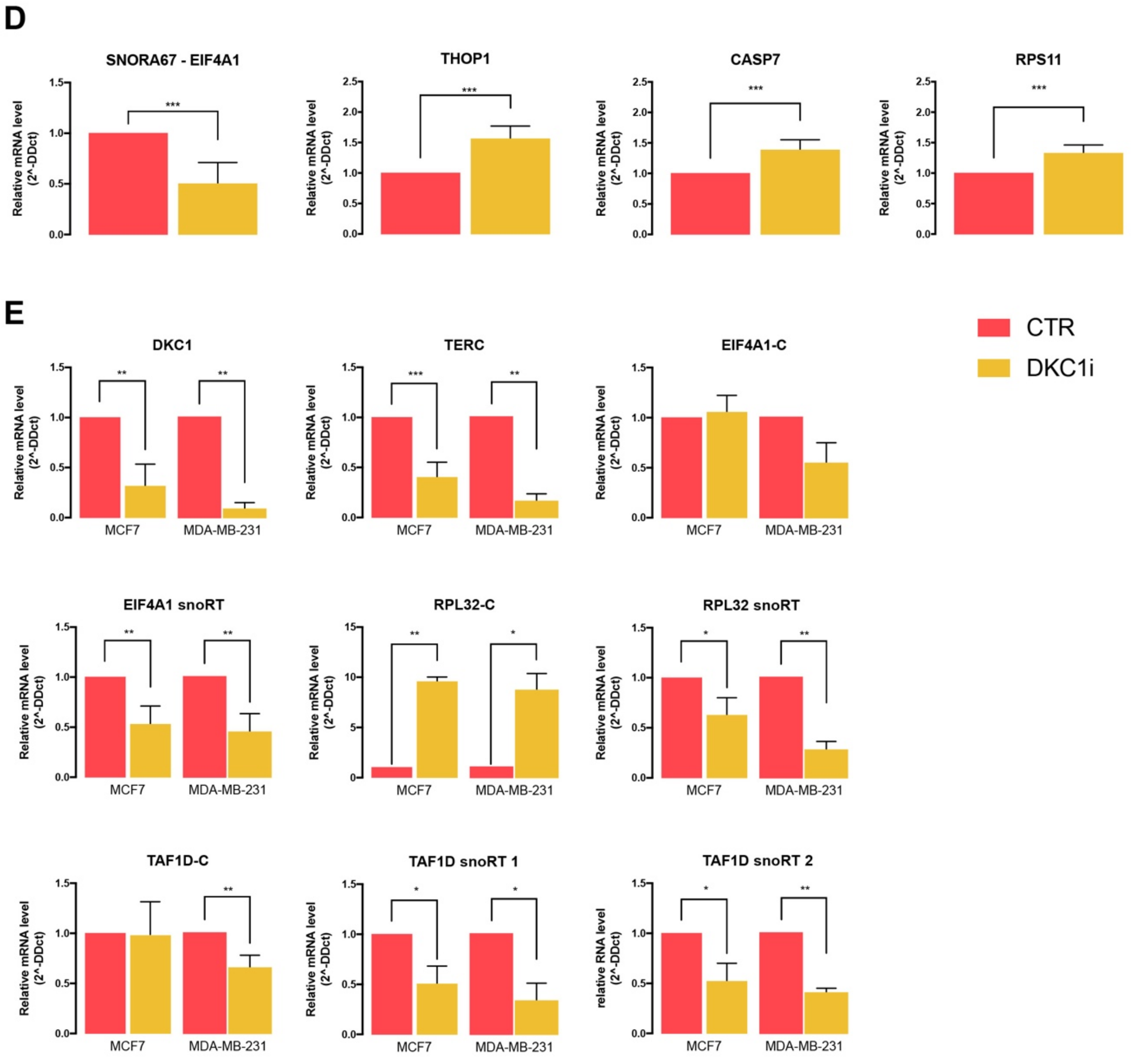

**Figure S1**, related to Figure 1. *Differential expression analysis of total cytoplasmic and polysomal recruited species and validation by RT-qPCR.*

Count-based differential expression analysis of total cytoplasmic (A), transcripts (B) (the same images as Figure 1C), and transcripts obtained from exon analysis (C). Data are reported for both total cytoplasmic and polysomal RNA analysis. Differentially expressed transcripts are depicted as red dots, while transcript gene names of interest are highlighted. Given the small number of differentially expressed transcripts in the polysomal transcript analysis, those are all individually labelled instead of being shown as red dots. Below every plot there is a table with all the corresponding regulated gene names. For the gene or transcript ID see Supplementary Tables S2 and S3 file. (D) Validation by RT-qPCR quantification of the specific transcripts differentially recruited on polysomes on the same MCF7 cytoplasmic cell lysate used for the RNA-seq. (E) RT-qPCR quantification of the transcripts found to be differentially recruited on the polysomes observed in MCF7 and MDA-MB 231 total cell lysates after dyskerin KD using siRNA. DKC1 and TERC are used for dyskerin silencing control. The means from three biological replicates (n = 3) are shown; error bars represent SD. Paired Student's t tests were performed on the controls. *p < 0.05, **p < 0.01, ***p < 0.005, ****p < 0.001.

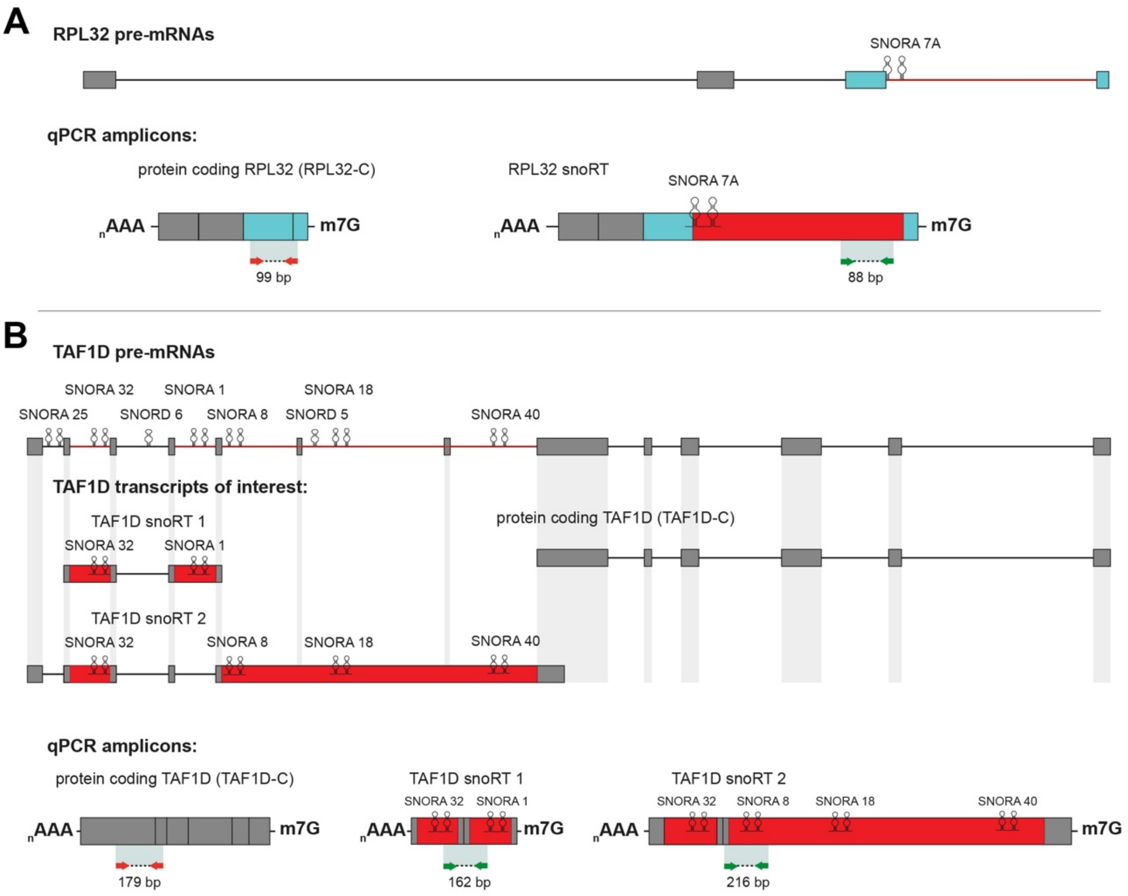

**Figure S2**, related to Figure 2. *Schematic overview of the intron-retaining isoforms of interest*.

(A) To-scale schematic overview of the *RPL32* pre-mRNA. Introns are depicted as lines connected to exons. *SNORA7A* is shown, and its intron is depicted as a red line in pre-mRNA or as a red box in RPL32 snoRT. *SNORA7A*-flanking exonic sequences are depicted as blue boxes. The qPCR amplicons are shown below the mRNA boxes, between arrows. The amplicon between red arrows identifies the protein coding mRNA, while the amplicon between green arrows identifies every RPL32 snoRT. (B) To-scale schematic overview of the *TAF1D* pre-mRNA. Introns are depicted as lines connected to exons. *SNORA25, SNORA32, SNORA1, SNORA8, SNORA18 SNORA40, SNORD6, and SNORD5* are shown. Intron-retaining SNORA sequences of interest are depicted either as a red line in pre-mRNA or as a red box in *TAF1D* transcripts of interest. qPCR amplicons are indicated. The amplicon between red arrows identifies the protein coding mRNA, while the amplicon between green arrows (different primers) specifically identifies the two transcripts shown in the figure. Primer sequences are listed in Supplementary Tables S1. m7G: cap; AAAn: poly(A) tail; P: monophosphate.

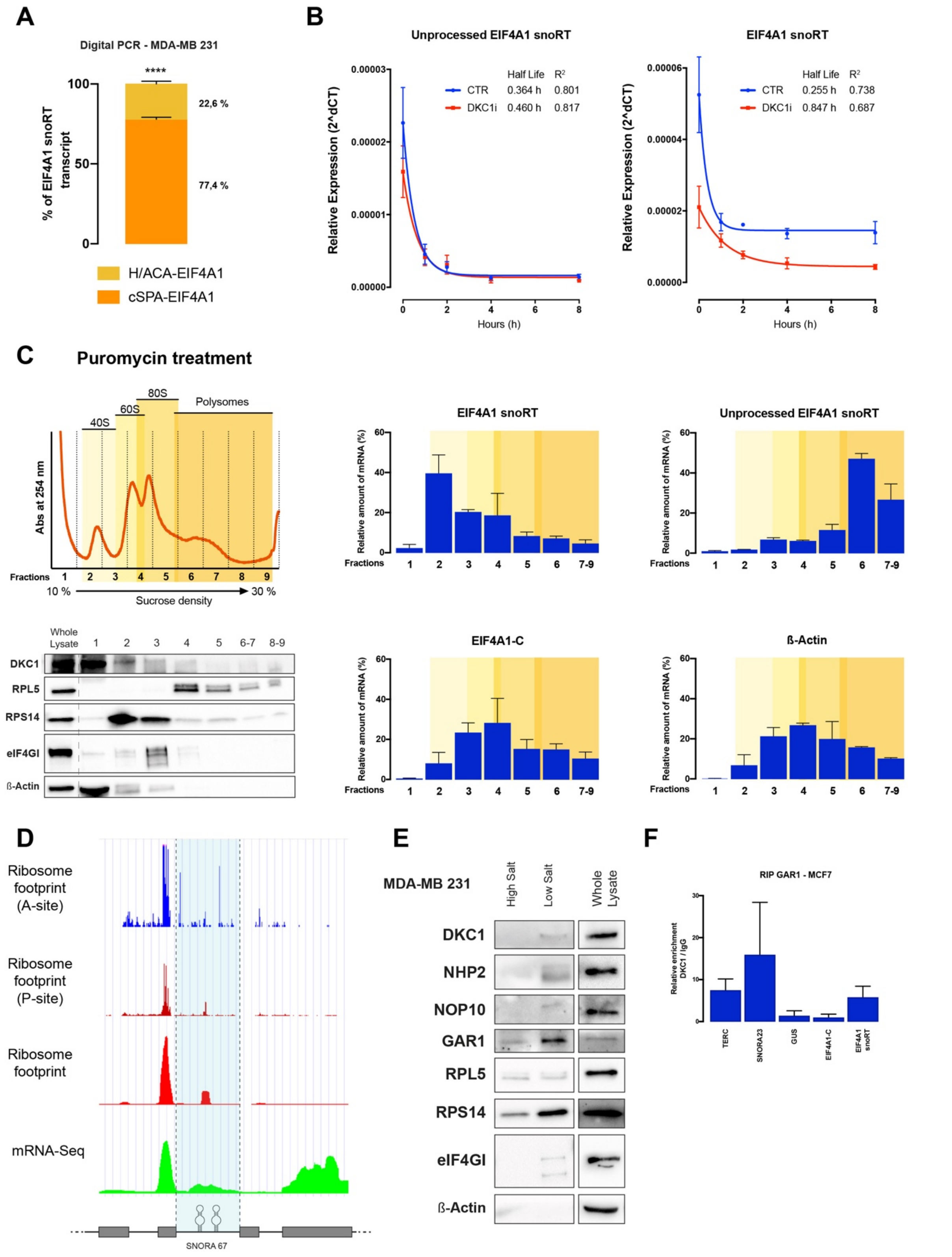

**Figure S3**, related to Figure 3. *Dyskerin-bound snoRT were processed to generate 5'snoRNA-ended polyadenylated transcripts and interact with ribosomes in the cytoplasm.*

(A) Percentage representation of EIF4A1 snoRT obtained by digital PCR absolute quantification in MDA-MB 231 cells. Data are shown as the percentage of EIF4A1 snoRT after normalization with GUS housekeeping transcript. The means from three biological replicates (n = 3) are displayed, error bars represent SD. Paired Student's t tests were performed on controls. *p < 0.05, **p < 0.01, ***p < 0.005, ****p < 0.001. (B) mRNA stability assay of EIF4A1 snoRT species after actinomycin D treatment. MCF7 cells with stable depletion of dyskerin and the relative controls were seeded at 70% confluence and incubated with actinomycin D (10 μg/ml) added to the medium. Cells were harvested at 0, 1, 2, 4, 8 h after treatment and RNA was extracted. Data are shown as relative expression (2^dCT) to the relatively stable rRNA 18S. The means from three biological replicates (n = 3) are shown, error bars represent SEM. (C) Polysome profiling analysis after puromycin treatment. Top left: representative polysome profile obtained by 10-30% sucrose density gradient centrifugation from MCF7 cells. Puromycin treatment prevents ribosome translocation during the elongation stage, resulting in a low number of polysomes obtained. The portions of the profile referring to the different ribosomal subunits are highlighted. Bottom left: distribution of dyskerin and control proteins across the gradient fractions analysed by Western blotting with specific antibodies. Right: Distribution of transcripts of interest after RNA purification from gradient fractions obtained by RT-qPCR. Results are expressed as the fraction (%) of the total amount of the transcripts contained in the lysate. Data are shown as means ± SEM of two different biological replicates. (D) Ribosome profile of global aggregate data obtained on UCSC Genome Browser database using GWIPS-vis tool. Only the portion near the SNORA 67 is shown. SNORA 67 intron is highlighted in blue. (E) Ribosome purification: Western blotting analysis of purified ribosomes from MDA-MB 231 cells shows a co-purification of all pseudouridine-RNP complex (DKC1, NHP2, NOP10, GAR1). RPL5 and RPS14 are shown as the positive control for the ribosomal purification, while eIF4G is used as the control for ribosome interacting factors. (F) RNA immunoprecipitation analysis of GAR1 from MCF7 total cellular lysates. RT-qPCR analysis of known dyskerin targets (TERC, SNORA23), a known off-target (GUS), and transcripts of interest. Results are expressed as the fold change against immunoprecipitation with IgG. Data are shown as means ± standard error of the mean (SEM).

**A**

| Transcripts biotype distribution | | |
|---|---|---|
| **Biotype** | **N°** | **%** |
| Protein Coding | 315 | 44,9 |
| Retained Intron | 125 | 17,8 |
| snoRNA / scaRNA | 115 | 16,4 |
| Processed Transcript | 61 | 8,7 |
| Nonsense Mediated Decay | 35 | 5,0 |
| lincRNA | 19 | 2,7 |
| Antisense | 18 | 2,5 |
| Other RNAs | 13 | 1,8 |

**B**

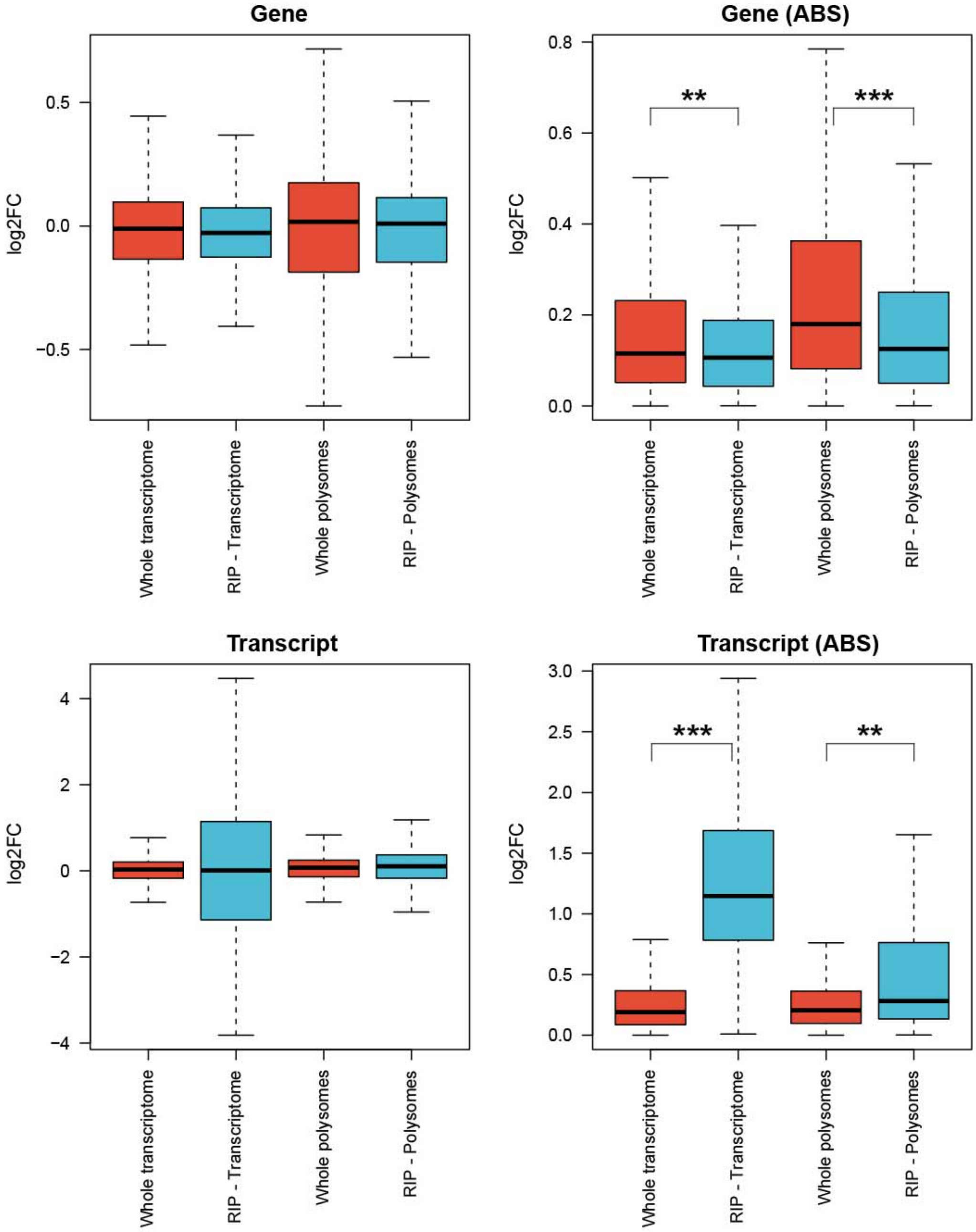

**C**

| Term | GO ID | Number of unique genes | Fold Enrichment | FDR *p*-value |
|---|---|---|---|---|
| **Gene Ontology: Molecular Function** | | | | |
| Nuclear hormone receptor binding | GO:0035257 | 11 | 6.14 | 1.99E-03 |
| RNA polymerase II-specific DNA-binding transcription factor binding | GO:0061629 | 19 | 5.82 | 4.81E-06 |
| Hormone receptor binding | GO:0051427 | 11 | 5.08 | 9.64E-03 |
| DNA-binding transcription factor binding | GO:0140297 | 21 | 5.02 | 5.38E-06 |
| Transferase activity, transferring acyl groups other than amino-acyl groups | GO:0016747 | 11 | 4.08 | 4.81E-02 |
| Transferase activity, transferring acyl groups | GO:0016746 | 12 | 3.92 | 4.07E-02 |
| Transcription factor binding | GO:0008134 | 24 | 3.02 | 1.54E-03 |
| Double-stranded DNA binding | GO:0003690 | 26 | 2.30 | 4.07E-02 |
| Protein binding | GO:0005515 | 185 | 1.35 | 6,15E-07 |
| Binding | GO:0005488 | 211 | 1.22 | 7.06E-06 |
| Molecular function | GO:0003674 | 226 | 1.13 | 7.37E-04 |
| **Gene Ontology: Biological Process** | | | | |
| Internal protein amino acid acetylation | GO:0006475 | 9 | 6.25 | 1.77E-02 |
| Internal peptidyl-lysine acetylation | GO:0018393 | 8 | 5.78 | 4.42E-02 |
| Protein acylation | GO:0043543 | 12 | 5.63 | 7.54E-03 |
| Protein acetylation | GO:0006473 | 9 | 5.44 | 2.81E-02 |
| Rhythmic process | GO:0048511 | 12 | 3.85 | 4.18E-02 |
| Peptidyl-amino acid modification | GO:0018193 | 26 | 2.61 | 1.62E-02 |
| Negative regulation of RNA metabolic process | GO:0051253 | 33 | 2.15 | 2.40E-02 |
| Positive regulation of transcription, DNA-templated | GO:0045893 | 36 | 2.04 | 2.40E-02 |
| Positive regulation of RNA metabolic process | GO:0051254 | 40 | 2.03 | 1.89E-02 |
| Positive regulation of nucleic acid-templated transcription | GO:1903508 | 38 | 2.03 | 2.16E-02 |
| Negative regulation of gene expression | GO:0010629 | 40 | 2.03 | 1.82E-02 |
| Positive regulation of RNA biosynthetic process | GO:0010630 | 38 | 2.03 | 2.10E-02 |
| Positive regulation of nucleobase-containing compound metabolic process | GO:0010631 | 41 | 1.91 | 3.87E-02 |
| Negative regulation of nitrogen compound metabolic process | GO:0010632 | 51 | 1.90 | 1.35E-02 |
| Negative regulation of cellular metabolic process | GO:0010633 | 53 | 1.82 | 1.82E-02 |
| Negative regulation of macromolecule metabolic process | GO:0010634 | 55 | 1.82 | 1.65E-02 |
| Negative regulation of metabolic process | GO:0010635 | 57 | 1.72 | 2.50E-02 |
| Positive regulation of nitrogen compound metabolic process | GO:0010636 | 62 | 1.71 | 1.69E-02 |
| Positive regulation of cellular metabolic process | GO:0010637 | 64 | 1.69 | 1.83E-02 |
| Cellular protein modification process | GO:0010638 | 59 | 1.69 | 2.51E-02 |
| Protein modification process | GO:0010639 | 59 | 1.69 | 2.43E-02 |
| Macromolecule modification | GO:0010640 | 62 | 1.66 | 2.36E-02 |
| Positive regulation of macromolecule metabolic process | GO:0010641 | 63 | 1.65 | 2.55E-02 |
| Positive regulation of metabolic process | GO:0010642 | 67 | 1.62 | 2.35E-02 |
| Cellular protein metabolic process | GO:0010643 | 68 | 1.62 | 2.31E-02 |
| Regulation of RNA metabolic process | GO:0010644 | 69 | 1.61 | 2.47E-02 |
| Cellular macromolecule metabolic process | GO:0010645 | 90 | 1.56 | 9.64E-03 |
| Regulation of gene expression | GO:0010646 | 79 | 1.56 | 1.84E-02 |
| Positive regulation of cellular process | GO:0010647 | 96 | 1.55 | 1.38E-02 |
| Negative regulation of cellular process | GO:0010648 | 83 | 1.54 | 1.84E-02 |
| Regulation of primary metabolic process | GO:0010649 | 104 | 1.52 | 1.82E-02 |
| Negative regulation of biological process | GO:0010650 | 91 | 1.51 | 1.74E-02 |
| Regulation of nitrogen compound metabolic process | GO:0010651 | 98 | 1.48 | 1.51E-02 |
| Positive regulation of biological process | GO:0010652 | 104 | 1.48 | 1.18E-02 |
| Regulation of metabolic process | GO:0010653 | 111 | 1.46 | 7.74E-03 |
| Regulation of cellular metabolic process | GO:0010654 | 103 | 1.46 | 1.40E-02 |
| Regulation of macromolecule metabolic process | GO:0010655 | 101 | 1.45 | 1.92E-02 |
| Cellular metabolic process | GO:0010656 | 118 | 1.36 | 2.36E-02 |
| Biological process | GO:0010657 | 226 | 1.12 | 8.92E-03 |
| **Gene Ontology: Cellular Component** | | | | |
| Nucleoplasm | GO:0005654 | 66 | 1.67 | 3.52E-03 |
| Nuclear part | GO:0044428 | 83 | 1.63 | 1.50E-03 |
| Nuclear lumen | GO:0031981 | 75 | 1.62 | 2.90E-03 |
| Organelle lumen | GO:0043233 | 84 | 1.41 | 5.64E-02 |
| Intracellular organelle lumen | GO:0070013 | 84 | 1.41 | 5.26E-02 |
| Membrane-enclosed lumen | GO:0031974 | 84 | 1.41 | 4.93E-02 |
| Nucleus | GO:0005634 | 113 | 1.35 | 1.95E-02 |
| Intracellular organelle part | GO:0044446 | 141 | 1.33 | 2.09E-03 |
| Organelle part | GO:0044422 | 145 | 1.33 | 1.98E-03 |
| Cytoplasmic part | GO:0044444 | 142 | 1.28 | 1.22E-02 |
| Intracellular membrane-bounded organelle | GO:0043231 | 159 | 1.28 | 2.63E-03 |
| Cytoplasm | GO:0005737 | 162 | 1.23 | 1.27E-02 |
| Intracellular organelle | GO:0043229 | 178 | 1.23 | 2.43E-03 |
| Intracellular part | GO:0044424 | 199 | 1.21 | 7.27E-04 |
| Intracellular | GO:0005622 | 199 | 1.20 | 1.45E-03 |
| Organelle | GO:0043226 | 183 | 1.19 | 1.20E-02 |
| Membrane-bounded organelle | GO:0043227 | 168 | 1.19 | 4.82E-02 |
| Cellular component | GO:0005575 | 228 | 1.07 | 5.25E-02 |
| **Gene Ontology: PANTHER Pathway** | | | | |
| CCKR signaling map | P06959 | 9 | 4.56 | 3.78E-02 |
| Gonadotropin-releasing hormone receptor pathway | P06664 | 10 | 3.84 | 3.25E-02 |

**Figure S4**, related to Figure 4. *Dyskerin binds to a complex RNA interactome in the cytoplasm and these genes are regulated after dyskerin partial depletion.*

(A) List of transcript biotype distribution and the number of genes identified by RIP-Seq analysis shown in Figure 4A. (B) Distribution of log2 (fold-change) in the shDKC1 vs CTRL MCF7 cells RNA-seq dataset at the transcriptome and polysome levels. These tables show the distribution for genes identified as differentially expressed in that dataset (red boxplots) and for genes identified as targets by the RIP-seq analysis (light blue boxplots), at the gene and transcript level (top and bottom row, respectively). Absolute fold change data are plotted on panels at right. (C) List of all statistically significant gene ontology terms broken down by molecular function, biological process, cellular component, and for the PANTHER pathways.

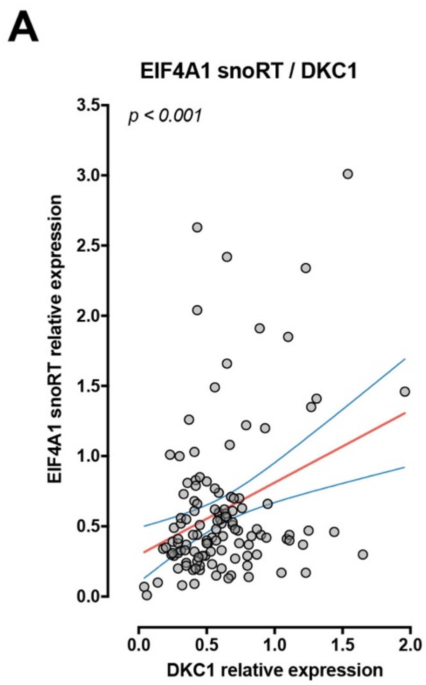

**Figure S5**, related to Figure 5.

(A) Correlation between DKC1 and EIF4A1 snoRT relative expression in RNA extracted from 120 breast cancer tissue samples. Data obtained by RT-qPCR. The blue lines represent 95% C.I., while the red line is the best-fitting line. Pearson correlation coefficient test was performed on controls.

**A**

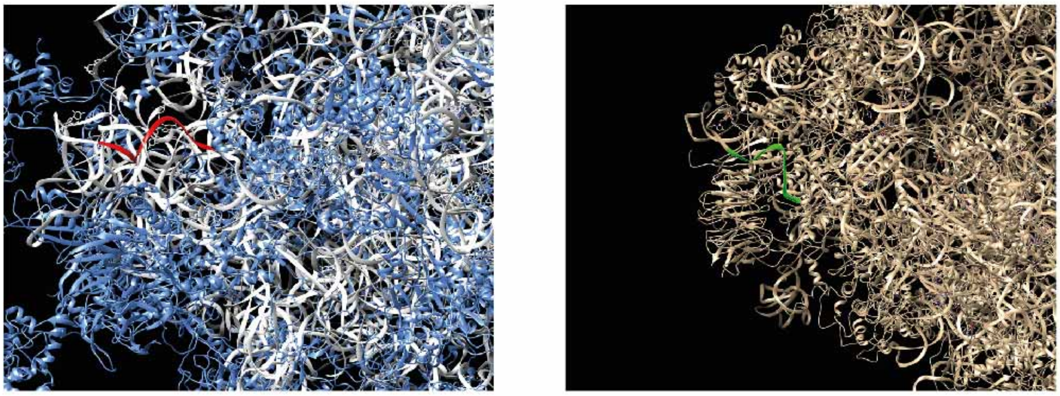

**Figure S6**. The SNORA67 target rRNA sequence is exposed on the ribosome.

3D structure of ribosome with the SNORA67 target sequence (which includes the U1445) highlighted. The left panel shows the target sequence in red; white ribbons are other rRNA sequences; blue ribbons are ribosomal proteins. The right panel shows the same structure from a different point of view. The target sequence is shown in green; other rRNA and protein sequences are shown in light brown.
